# Aberrant regulation of the Rap1 small GTPase in response to escalating, intermittent stress produces hippocampal synaptic and cognitive dysfunction

**DOI:** 10.1101/2023.07.03.547282

**Authors:** Kathryn J. Bjornson, Amanda M. Vanderplow, Yezi Yang, Bailey A. Kermath, Michael E. Cahill

**Author notes:** Corresponding author Address: 2015 Linden Drive, Madison, WI, 53706, Contact.

## Abstract

The effects of repeated stress on cognitive impairment are thought to be mediated, at least in part, by reductions in the stability of dendritic spines in brain regions critical for proper learning and memory, including the hippocampus. Small GTPases are particularly potent regulators of dendritic spine formation, stability, and morphology in hippocampal neurons. Through the use of small GTPase protein profiling in mice, we identify increased levels of synaptic Rap1 in the hippocampal CA3 region in response to escalating, intermittent stress. We then demonstrate that increased Rap1 in the CA3 is sufficient in and of itself to produce stress-relevant dendritic spine and cognitive phenotypes. Further, using super-resolution imaging, we investigate how the pattern of Rap1 trafficking to synapses likely underlies its effects on the stability of select dendritic spine subtypes. These findings illuminate the involvement of aberrant Rap1 regulation in the hippocampus in contributing to the psychobiological effects of stress.

## Introduction

Stress is thought to be a major precipitating factor for the onset of several psychiatric disorders.^1–9^ In addition, stress is a distinct psychobiological process unto itself that can produce a well-characterized series of physical consequences.^10–17^ Stress also can elicit a constellation of behavioral effects including altered executive and cognitive function, sleep alterations, and changes in mood.^18–21^

In rodents, multiple forms of severe stress have been shown to alter the stability of dendritic spines in pyramidal neurons of the medial prefrontal cortex (mPFC) and hippocampus.^22–30^ Within the hippocampus, the synaptic density along CA3 pyramidal neurons is commonly afflicted by stress.^24,26^ Although some forms of stress exposure can affect CA1 dendritic spine density,^23,30^ there are several examples of stress affecting dendritic spine stability in CA3 neurons with no effect in CA1 neurons.^24,26^ Interventions that alleviate the effects of stress on hippocampal neuronal structure are frequently associated with a mitigation of stress-driven behavioral aberrations.^31,32^

Human studies also suggest that stress has detrimental consequences on the CA3 region. Notably, heightened levels of perceived stress in adult humans are associated with reduced CA2/CA3 volume with no corresponding effect on CA1 volume or on whole brain volume.^33^ Further, elevated levels of trait anxiety in adult humans are associated with a significant reduction in CA3 dendritic spine density.^34^ In addition, elevated cortisol levels in children are associated with significant reductions in CA3 and dentate gyrus volume, yet no change in CA1 volume.^35^

Single bout (acute) restraint and daily, multi-hour restraint that spans a week or more (chronic) have emerged as a common means of inducing a form of severe, inescapable stress in rodents.^26,29,36,37^ We recently developed and performed an initial characterization of an escalating, intermittent restraint stress protocol in mice that administers 6 days of restraint stress across an 8 day period.^38^ This escalating, intermittent stress protocol was conceptualized as a way to model the notion that stress severity is not always uniform and does not always occur on successive days, while simultaneously recognizing that minor stressors can transition to major stressors in a short duration. Our prior findings indicate that this stress protocol is capable of producing striking cognitive impairment, including deficits in spatial working memory and object-in-place memory. In addition, this escalating, intermittent stress protocol causes a reduction in dendritic spine density in mPFC pyramidal neurons that closely mirrors the magnitude of spine loss seen with longer multi-week stress administration.^38^ However, the effects of escalating, intermittent stress on dendritic spine phenotypes in hippocampal neurons, including those of the CA3 field, is completely unknown.

Here we examine the effects of escalating, intermittent stress in mice on the dendritic spine density and morphology of hippocampal pyramidal neurons. Our findings indicate that this stress procedure causes a striking reduction in spine stability in CA3 pyramidal neurons that has some important differences compared to what we recently reported in the mPFC. Members of the Ras superfamily of small GTPases act as molecular switches that are highly enriched in many neuronal regions, including dendritic spines, where they potently impact dendritic spine formation, stability, and maturation.^39^ Here we use subcellular fractionation to profile multiple major small GTPases known to be enriched in hippocampal pyramidal neurons. Our findings identify the specific upregulation of the Rap1 small GTPase in synaptic fractions of the CA3 hippocampal region in response to escalating, intermittent stress.

Like many small GTPases, Rap1 is enriched in dendritic spines, and its activation is largely controlled by guanine nucleotide exchange factors (GEFs) that induce a GDP to GTP exchange on small GTPases, thereby facilitating their activation.^39^ Several Rap1-specific GEFs have been identified, many of which are also localized to dendritic spines where they facilitate dendritic spine destabilization in a manner requiring Rap1 engagement.^40^ In addition, Rap1 directly interacts with multiple PDZ domain-containing scaffolding proteins that are enriched in dendritic spines, including SHANK3 ^41^ and AF6 (afadin).^42,43^

At present, the majority of neural studies of Rap1’s function have examined cultured pyramidal neurons with a scarcity of *in vivo* assessments. Here we demonstrate that increased Rap1 in the CA3 *in vivo*, as occurs in response to escalating, intermittent stress, is sufficient to produce stress-relevant synaptic phenotypes, namely a reduced density of thin dendritic spines, in addition to stress-relevant cognitive impairments. Finally, we demonstrate that the pattern of Rap1 trafficking to dendritic spines potentially explains the particularly striking effects of Rap1 on the destabilization of thin dendritic spines. Taken together with our prior study in mPFC neurons,^38^ these hippocampal findings implicate a multi-forebrain region involvement of Rap1 in potentially mediating the detrimental effects of escalating, intermittent stress on synaptic stability and cognition.

## RESULTS

### Escalating, intermittent stress alters c-Fos levels in the CA3 region of the hippocampus

Increased levels of immediate early genes (IEGs), such as *c-Fos*, commonly accompany neuronal activation. As such, fluctuations in the expression of IEGs following exposure to novelty or in response to cognitive processing are a commonly used and highly quantifiable indicator of recent neuronal activation.^44–46^ Prior studies using the administration of a single bout of stress have mapped changes in c-Fos mRNA and protein across many forebrain regions, including the hippocampus. However, within the hippocampus, including the CA3 field, the findings from these single stress exposure studies have been largely inconsistent, ranging from a lack of identified changes in IEG expression to significant c-Fos induction.^47,48^ Complicating matters further, the effects of repeated stress exposure on c-Fos levels within the forebrain has received comparatively minimal attention, with no studies examining escalating, intermittent stress.

We previously showed that c-Fos transcript is rapidly upregulated in the medial prefrontal cortex of mice following the free exploration of novel objects.^38,49^ Here we wanted to determine if novel object exploration increases c-Fos protein levels in the CA3, and if so, whether prior exposure to escalating, intermittent stress impacts any observed changes in c-Fos induction. To this end, we exposed young adult mice to escalating, intermittent stress using our established procedures in which mice are subjected to 6 days of variable duration physical restraint over an 8 day period. 24 hours following the final stress exposure, mice were randomly assigned to either an object exploration condition or to a cage control condition. Non-stressed mice were used for baseline comparison. Mice in the object exploration condition were allowed to explore four novel objects in an open arena for 7 minutes, and were transcardially perfused 3 hours later (**Figure 1A**). Cage control mice were never exposed to the novel objects. Importantly, our stress protocol does not affect the amount of time mice spend exploring objects [t(8)=1.328, p=0.2209] (**Figure 1B**), and thus escalating, intermittent stress does not impact novelty-seeking motivation. Using immunofluorescence, we quantified the number of c-Fos expressing cells in both hemispheres of the CA3 from three coronal plane sections. The coronal section rostral/caudal levels used for analysis were equivalent across all mice, thereby allowing for direct comparisons between different animals. For statistical analysis, each mouse (rather than each brain section) was used as an n.

**Figure 1.**
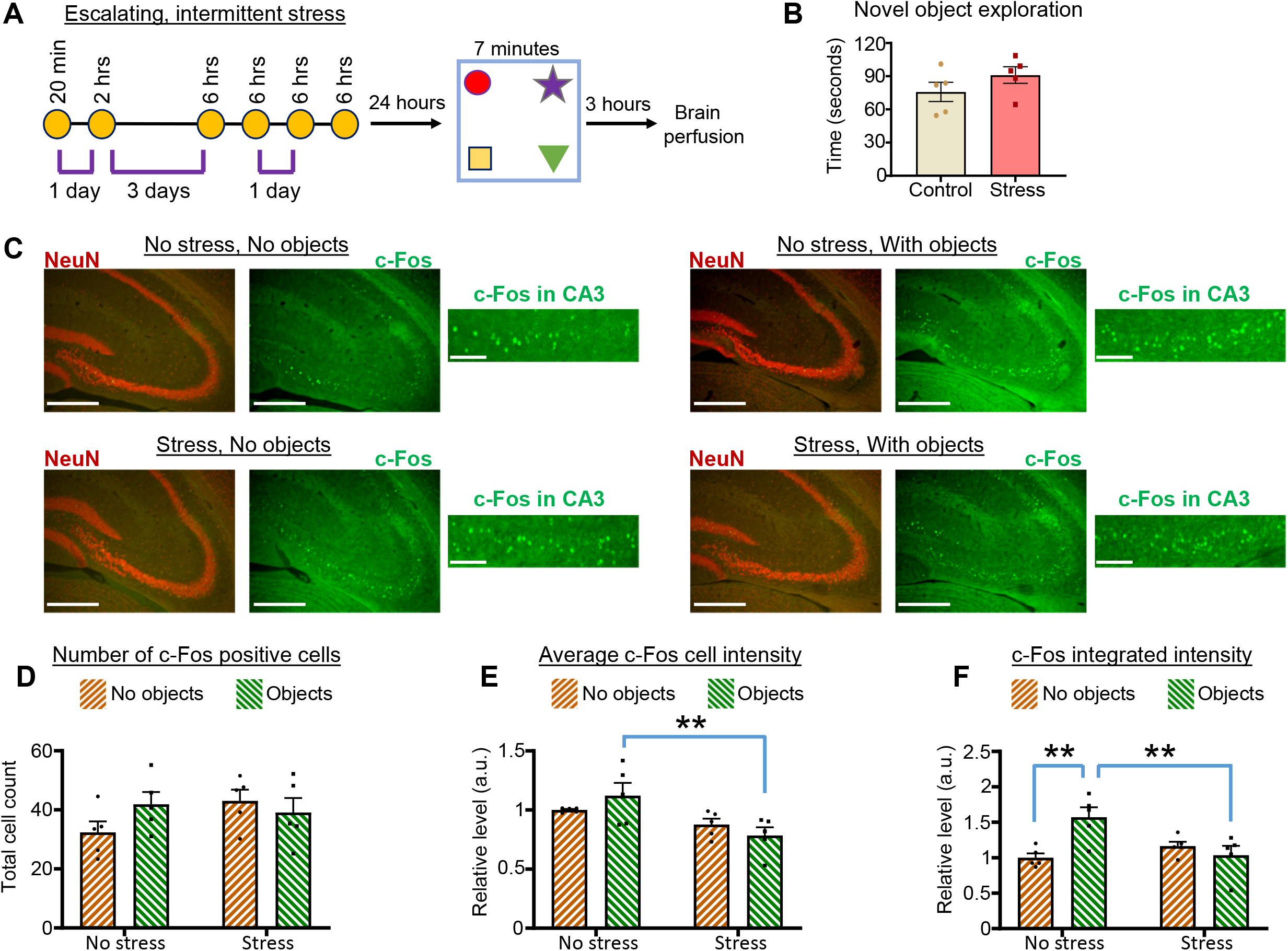
Escalating, intermittent stress increases c-Fos immuno-reactivity in the CA3 region. A. Schematic of the escalating, intermittent stress protocol procedure. On day 1, mice were restrained for 20 minutes, followed by 2 hours of restraint on day 2. Mice then had two consecutive days with no restraint. This was followed by four consecutive days of 6 hour restraint. 24 hours following the final day of restraint, mice were allowed to explore four novel objects in an open field for 7 minutes. Mice were then euthanized 3 hours after the object exploration. Mice that were not exposed to novel objects were used as a control. B. Graph depicts the amount of time non-stressed mice and mice exposed to escalating, intermittent stress spent investigating novel objects during a 7 minute period. Stress mice showed a normal motivation to explore novel objects. n=5 no stress mice, 5 stress mice. C. Low magnification images show NeuN and c-Fos immuno-label in the hippocampus (with CA3 region visible) of non-stressed and stressed mice that were either not exposed to novel objects or were allowed to explore novel objects (Scale bar=300µm). High magnification images exclusively show the CA3 region and the associated c-Fos immuno-label (Scale bar=10µm). D. Graph depicts the number of c-Fos positive cells in the CA3 of each stress and object exploration condition. No main effect or differences between groups were detected. Total cell count is the combined total from three coronal planes of the CA3. n=5 no stress, no object mice; 5 no stress, object mice; 5 stress, no object mice; 5 stress, object mice. E. Graph depicts the average c-Fos immunofluorescent intensity among CA3 region c-Fos positive cells in each stress and object exploration condition. The c-Fos intensity of all cells within a single mouse were averaged, and this value used for statistical analyses. **Bonferroni, p<0.01. n is same as indicated in 1D. F. Graph depicts c-Fos integrated intensity in each stress and object exploration condition. The integrated intensity of all cells within a single mouse were averaged, and this value used for statistical analyses. **Bonferroni, p<0.01. n is same as indicated in 1D. All summary data are the mean + SEM.

Surprisingly, we found that neither stress nor object investigation affected the number of c-Fos positive cells in the CA3 [stress main effect, F(1,16)=0.8837, p=0.3612; object investigation main effect, F(1,16)=0.4447, p=0.5144; all Bonferroni post hoc comparisons p>0.05] (**Figures 1C and 1D and Figure S1A**). Further, no differences in the density of c-Fos positive cells in the CA3 were detected (cells per area of the CA3) (**Figure S1B**). Within the population of c-Fos positive cells, the relative amount of c-Fos expressed is an important factor. We thus quantified the fluorescent intensity of the c-Fos fluorescent label in all c-Fos positive cells in the CA3. Among the population of c-Fos positive cells, we detected a main effect of stress status, but no main effect of object investigation status and no interaction, for the average c-Fos intensity [stress main effect, F(1,16)=11.06, p=0.0043; object investigation main effect, F(1,16)=0.04281, p=0.8387; interaction, F(1,16)=2.401, p=0.1408]. Further, post hoc testing within non-stressed and stressed mice did not reveal an effect of object investigation on c-Fos average label intensity (Bonferroni post hoc: non-stressed object vs. non-stressed no object, p=0.4643; stressed object vs. stressed no object, p=0.7131). However, post hoc testing revealed a significant increase in the average c-Fos label intensity in the non-stressed object investigation group vs. the stressed object investigation group (Bonferroni post hoc, p=0.0066) (**Figures 1C and 1E**).

As an indicator of total c-Fos levels in the CA3, we examined the integrated intensity of the c-Fos label within individual mice, defined as the product of the number of c-Fos expressing cells and the average c-Fos fluorescent intensity of individual cells. For integrated intensity, we identified a significant interaction between stress status and object investigation status [F(1,16)=11.04, p=0.0043]. Among non-stressed mice, post hoc testing revealed a significant increase in c-Fos integrated intensity in mice that explored objects as compared to cage control mice (Bonferroni post hoc, p=0.0029). However, no such increase in c-Fos integrated intensity in response to object exploration was identified in mice exposed to stress (Bonferroni post hoc, p=0.8041). Further, among object exploration mice, the c-Fos integrated intensity was significantly greater for non-stressed mice as compared to stressed mice (Bonferroni post hoc, p=0.0049) (**Figures 1C and 1F**). Similar findings were identified when the integrated intensity was corrected for the measured area of the CA3 field (**Figure S1C**). The elevated c-Fos integrated intensity in the CA3 of non-stressed mice that experienced object exploration is likely due to compounding effects of non-significant increases in both the number of c-Fos positive cells and the average c-Fos fluorescent intensity. These findings suggest that escalating, intermittent stress attenuates c-Fos induction within the CA3 in response to novelty exploration.

### Escalating, intermittent stress impacts pyramidal neuron dendritic spine stability in the CA3 region

Escalating, intermittent stress produces synapse destabilization in mPFC pyramidal neurons.^38^ However, the effects of this stress procedure on dendritic spine density and morphology in hippocampal pyramidal neurons remains unknown. To this end, we subjected young adult mice to our escalating, intermittent stress procedure of 6 day variable duration physical restraint across an 8 day period, and 24 hours following the final stress exposure, mice were transcardially perfused (**Figure 2A**). Non-stressed mice were used as a control group. In *ex vivo* coronal brain sections, DiOlistic delivery of DiI red fluorescence (1,1’-dioctadecyl-3,3,3’3’- tetramethylindocarbocyanine perchlorate) was used to illuminate random populations of hippocampal neurons (**Figure 2B**). We focused our analysis on neurons residing within the CA3 field of the hippocampus. To reduce baseline variability in spine density caused by dendrite location, we exclusively imaged dendritic spines located on secondary dendrites of the apical tree,^50,51^ as done in prior hippocampal stress studies by other groups.^22,24^

**Figure 2.**
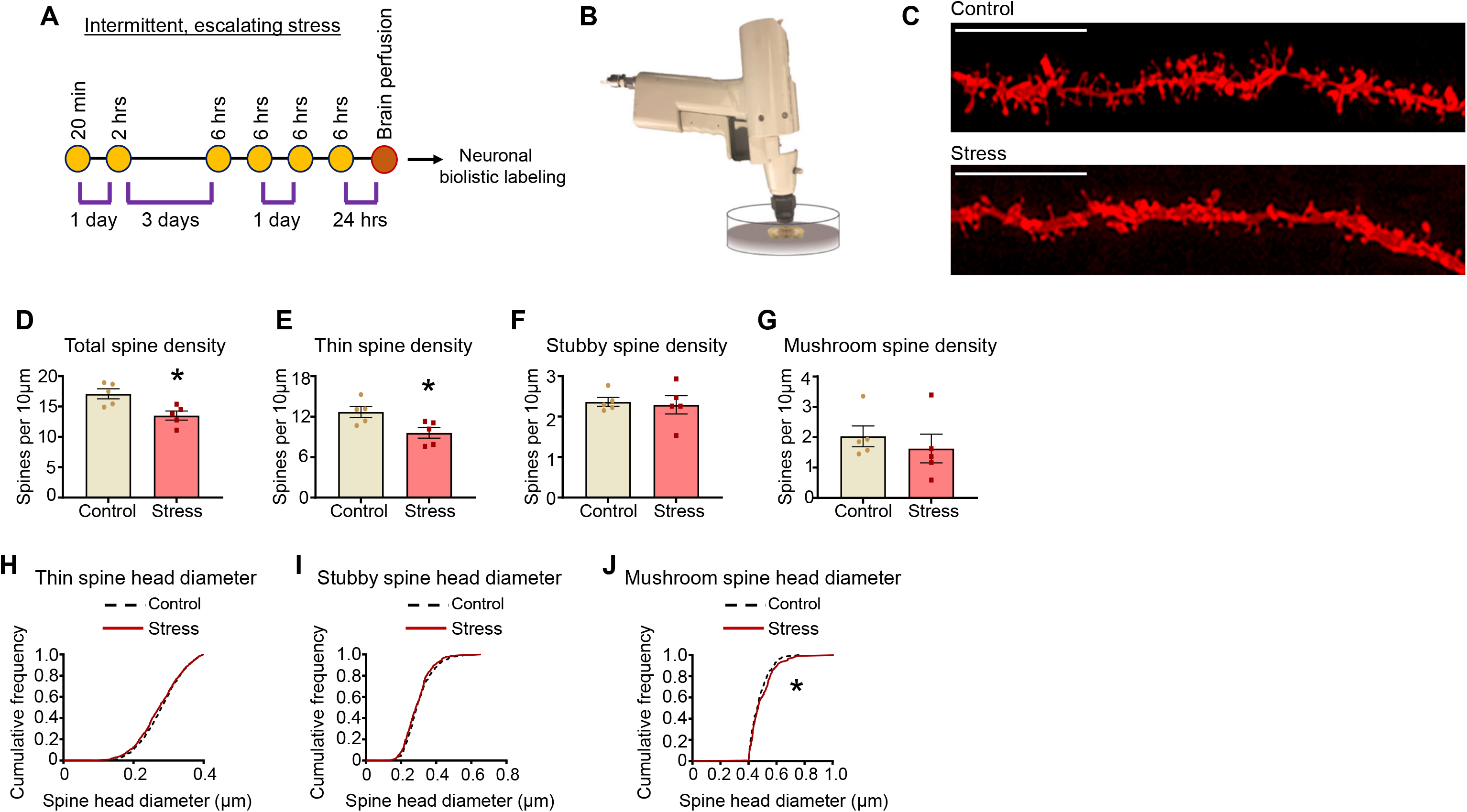
Escalating, intermittent stress reduces dendritic spine stability in CA3 pyramidal neurons. A. Schematic of experimental design. Mice were transcardially perfused 24 hours following the final stress exposure. B. Coronal mouse brain sections containing the dorsal hippocampus were labeled with DiI using a helium pressurized gene gun. C. Representative images of CA3 region pyramidal neuron secondary dendrites of the apical tree from non-stress and stressed mice. Scale bar=10µm D. Graph depicts the total dendritic spine density in non-stressed and stressed mice. *Two-tailed t-test, p<0.05. 3-5 neurons were imaged per mouse and spine values across all neurons within a single mouse were averaged. n=5 non-stress and 5 stress mice. E. Graph depicts thin dendritic spine density in non-stressed and stressed mice. *Two-tailed t-test, p<0.05. 3-5 neurons were imaged per mouse and spine values across all neurons within a single mouse were averaged. n=5 non-stress and 5 stress mice. F. Graph depicts stubby spine density in non-stressed and stressed mice. No significant differences were identified. 3-5 neurons were imaged per mouse and spine values across all neurons within a single mouse were averaged. n=5 non-stress and 5 stress mice. G. Graph depicts mushroom spine density in non-stressed and stressed mice. No significant differences were identified. 3-5 neurons were imaged per mouse and spine values across all neurons within a single mouse were averaged. n=5 non-stress and 5 stress mice. H. No significant differences in the cumulative thin spine head diameter curve were identified between the non-stress and stress conditions. n=1360 spines in non-stressed mice and 1231 spines in stressed mice. I. No significant differences in the cumulative stubby spine head diameter curve were identified between the non-stress and stress conditions. n=251 spines in non-stressed mice and 274 spines in stressed mice. J. A modest, yet significant rightward shift in the cumulative mushroom spine head diameter curve was identified in stressed mice vs. non-stressed mice. *Mantel-Cox, p<0.05. n=221 spines in non-stressed mice and 199 spines in stressed mice. All summary data are the mean + SEM.

Dendritic spine density and the head diameter of individual spines were semi-automatically quantified using the NeuronStudio program as detailed in our previous work.^38,49,52^ Within individual mice, the spine density for all neurons was averaged to a single data point, such that the n for statistical analysis is the number of mice assessed in the control and stress groups, rather than the total number of neurons. We found that escalating, intermittent restraint stress reduced the total density of dendritic spines in CA3 pyramidal neurons [t(8)=3.233, p=0.0120] (**Figures 2C and 2D**). To determine whether stress affects a particular subclass of dendritic spines, spines were categorized as thin, stubby, or mushroom based on our established criteria.^38,49,52^ Briefly, stubby spines are those that lack a discernable head, whereas thin and mushroom spines have a clear neck region. Neck-bearing spines with a head diameter of 0.4µm or greater were classified as mushroom, while neck-bearing spines with a head diameter below this value were categorized as thin. Spine subtype analysis revealed that escalating, intermittent stress reduced the density of thin dendritic spines in CA3 pyramidal neurons [t(8)=2.766, p=0.0244] (**Figure 2E**). Conversely, stress had no impact on the density of stubby [t(8)=0.2949, p=0.7755] or mushroom spines [t(8)=0.6930, p=0.5079] (**Figures 2F and 2G**).

Within the spine subtypes of thin, stubby, and mushroom, individual spines have a range of head diameters. We thus evaluated whether there were any changes in head diameter frequencies within spine subtypes of stress vs. control group neurons. To this end, we assessed cumulative spine head diameter frequencies, which can reveal subtle changes in subtype head diameter populations that are often missed when considering mean values alone. We found no significant differences in the thin and stubby spine head cumulative frequency curves between the stress and control groups (thin, χ2=1.524, p=0.2171; stubby, χ2=1.476, p=0.2244) (**Figures 2H and 2I**). On the other hand, we detected a subtle, yet significant, rightward shift in the spine head cumulative frequency curve for mushroom spines in the stress group relative to controls (χ2=4.469, p=0.0345) (**Figure 2J**). Given that stress did not alter mushroom spine density, the slight rightward shift in the mushroom head cumulative frequency curve is not likely due to the *de novo* formation of a cluster of mushroom spines containing unusually large heads. Rather, these data suggest that stress caused a subtle increase in head diameter within a population of pre-existing mushroom spines.

### Escalating, intermittent stress impacts surface GluA1 levels in the CA3 region

Surface AMPA receptors (AMPARs) on dendritic spines mediate many key aspects of the excitability of neurons.^53^ To determine if the reduction in CA3 dendritic spine density resulting from escalating, intermittent stress is associated with alterations in the abundance of surface AMPA receptors, we used Bis(sulfosuccinimidyl) suberate (BS^3^) crosslinking to examine surface and internal levels of the GluA1 AMPA receptor subunit^54–56^ (**Figure 3A**). We found that escalating, intermittent stress had no impact on total GluA1 levels [t(10.4)=0.2231, p=0.8277] (**Figures 3B and 3C**), yet stress decreased surface to internal ratios of GluA1 [t(10.09)=2.515, p=0.0305] (**Figures 3B and 3D**). Further analysis indicates that this decreased ratio in stressed mice is due to a decrease in surface GluA1 [t(8.735)=2.467, p=0.0365] (**Figures 3B and 3E**) and a proportional increase in levels of internal GluA1 [t(9.592)=2.406, p=0.0379] (**Figures 3B and 3F**). As the GluA1 subunit confers increased calcium permeability to AMPA receptors,^57,58^ reduced levels of surface GluA1 in stressed mice are suggestive of reduced AMPAR function.

**Figure 3.**
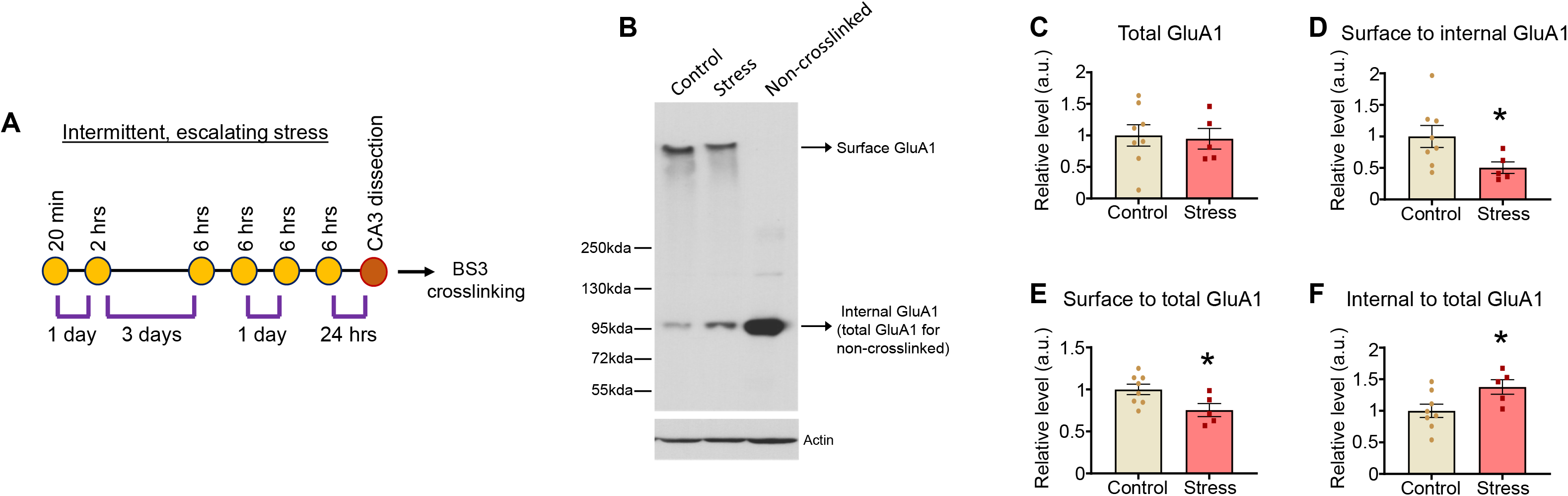
Escalating, intermittent stress reduces surface GluA1 levels in the CA3 region. A. Schematic of experimental design. CA3 region was microdissected 24 hours following the final stress exposure and prepared for BS^3^ crosslinking. B. Representative Western blot showing surface and internal GluA1 levels in the CA3 region of non-stressed (control) and stress mice. A non-crosslinked sample was used as a negative control for surface GluA1. C. Quantification of total GluA1 levels (surface+internal) in non-stressed and stressed mice. No differences between groups were observed. n=8 control mice, 5 stress mice D. Quantification of surface GluA1 to internal GluA1 ratios identified a significant decrease in stress mice compared to control mice. *Two-tailed t-test, p<0.05. n=8 control mice, 5 stress mice E. Quantification of surface GluA1 levels normalized to total GluA1 levels. A significant decrease in surface GluA1 was detected in stress mice. *Two-tailed t-test, p<0.05. n=8 control mice, 5 stress mice F. Quantification of internal GluA1 levels normalized to total GluA1 levels. A significant increase in internal GluA1 was detected in stress mice. *Two-tailed t-test, p<0.05. n=8 control mice, 5 stress mice All summary data are the mean + SEM.

### The effects of escalating, intermittent stress on small GTPase expression profiles in the CA3 region

The mechanisms that mediate the effects of stress on synaptic destabilization are undoubtedly complex and likely involve a constellation of molecular and biochemical alterations. Due to their potent ability to cause actin cytoskeletal rearrangements within synapses, there has been much interest in investigating the potential role of small GTPases in stress responses, including their involvement in stress-mediated changes in dendritic spine stability. To determine if escalating, intermittent stress alters the expression profiles of small GTPases in the CA3, we microdissected the CA3 region from young adult male mice 24 hours following the final stress exposure of the escalating, intermittent stress procedure. Tissue was collected from three independent experiments to allow for a rigorous identification of any consistent changes, with a total n of 15 control and 16 stress mice. In addition to synapses, small GTPases are trafficked to many other cellular regions including the nucleus and throughout the cytoplasm.^39,59–62^ To examine small GTPase expression levels in multiple cellular compartments, CA3 homogenates were subcellularly fractionated into P1 (crude nuclear), S2 (cytosolic) and P2 (crude synaptosome) fractions using our previously validated procedures.^38^ Fractions were resolved via SDS-PAGE with subsequent Western blotting to probe for the Rap1, Ras, Cdc42, and Rac1 small GTPases. In P2 fractions, we identified a significant increase in levels of Rap1 (Rap1b isoform; Rap1a isoform was subthreshold in P2 fractions) in the stress group relative to non-stressed controls (Bonferroni p=0.0318) (**Figures 4A-4C and Figure S2A and S2B**). Importantly, within all three experiments, the stress group showed a substantial increase in P2 Rap1 levels relative to the control group of the same experiment (58%, 42%, and 71% increase), indicating that this effect was highly consistent and reproducible (**Figures S2C-S2F**). In contrast, no P2 fraction changes were identified in any of the other examined small GTPases (Ras Bonferroni p>0.9999; Rac1 Bonferroni p=0.7829; Cdc42 Bonferroni p=0.4238) (**Figures 4A-4C and Figures S2B**). Further, no changes in any of the examined small GTPases, including Rap1, were identified in the P1 fraction (Rap1 Bonferroni p>0.9999, Ras Bonferroni p>0.9999; Rac1 Bonferroni p>0.9999; Cdc42 Bonferroni p=0.7501) (**Figures 4D-4F**). Similarly, stress had no effect on levels of any examined small GTPases in the S2 fraction (Bonferroni p>0.9999 for Rap1, Ras, Rac1, and Cdc42) (**Figures 4G-4I**).

**Figure 4.**
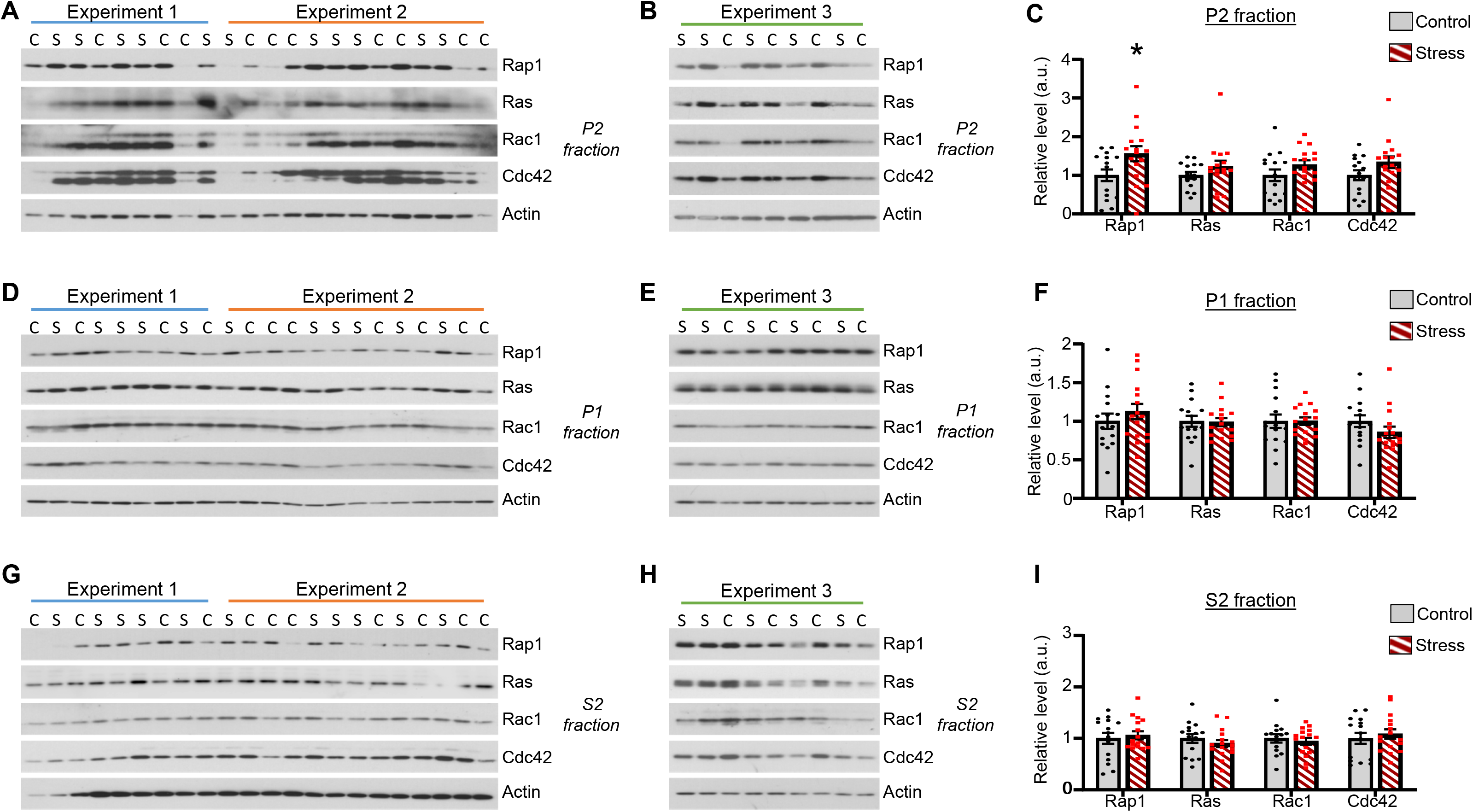
Escalating, intermittent stress increases Rap1 levels in the P2 fraction of CA3 region homogenates. A and B. P2 fractions derived from CA3 homogenates of non-stressed mice and mice subjected to escalating, intermittent stress (tissue collected 24 hour after final stress exposure). Western blots were probed for the indicated small GTPases. Samples were derived from 3 independent stress experiments. C=non-stressed controls, S=stressed. C. Stress increased Rap1 in the P2 fraction, with no effect on the other small GTPases. *Bonferroni, p<0.05. n=15 non-stressed and 16 stressed mice. Direct comparison p values (Fisher’s LSD): Rap1 p=0.0079; Ras p=0.2799; Rac1 p=0.1957; Cdc42 p=0.1060. D and E. P1 fractions derived from CA3 homogenates of non-stressed mice and stressed mice. Blots were probed for the indicated small GTPases. Samples were derived from 3 independent stress experiments. C=non-stressed controls, S=stressed. F. Lack of effect of stress on the levels of any small GTPases in P1 fractions. n=15 non-stressed and 16 stressed mice. Direct comparison p values (Fisher’s LSD): Rap1 p=0.2576; Ras p=0.9073; Rac1 p=0.9904; Cdc42 p=0.1965. G and H. S2 fractions derived from CA3 homogenates of non-stressed mice and stressed mice. Blots were probed for the indicated small GTPases. Sample were derived from 3 independent stress experiments. C=non-stressed controls, S=stressed. I. Lack of effect of stress on the levels of any small GTPases in S2 fractions. n=15 non-stressed and 16 stressed mice. Direct comparison p values (Fisher’s LSD): Rap1 p=0.6271; Ras p=0.4169; Rac1 p=0.6289; Cdc42 p=0.4973. All summary data are the mean + SEM.

The prior biochemical analyses were all performed using male mice. As such, we wanted to determine if stress also impacts Rap1 levels in CA3 P2 fractions of female mice and/or alters the expression profile of other major small GTPases is any subcellular fractions in female mice. To this end, we subjected female mice to escalating, intermittent stress and micro-dissected the CA3 region 24 hours later. Interestingly, we found no significant effects of stress on the levels of Rap1, Ras, Rac1, or Cdc42 in CA3 P1, S2, or P2 fractions (**Figures S3A-S3F**). These findings indicate that the effects of escalating, intermittent stress on increased P2 fraction Rap1 levels are sex-dependent.

To determine if stress-mediated increases in hippocampal P2 fraction Rap1 in male mice is specific to the CA3 region or is common to other areas of the hippocampus, we also examined levels in the CA1 region and dentate gyrus. We found that escalating, intermittent stress had no effect on P2 fraction Rap1 levels in the CA1 region or dentate gyrus of male mice (**Figure S4A-S4D**). These findings indicate that the effects of stress on increased P2 fraction Rap1 are specific to the CA3 region and are not common to other examined hippocampal areas.

### Rap1 alters the stability of dendritic spine subtypes in CA3 region pyramidal neurons

Our biochemical profiling studies identify a selective stress-mediated increase in P2 fraction Rap1 with no corresponding changes in Rac1, Cdc42, or Ras. Rap1 is highly enriched in dendritic spines.^63^ To determine if Rap1 overexpression in CA3 neurons is sufficient to cause spine alterations that resemble those of escalating, intermittent stress, we used herpes simplex virus (HSV) to co-overexpress Rap1 (Rap1b isoform) and green fluorescent protein (HSV-Rap1-GFP). HSVs only infect neurons, with no effects on glia,^64^ and we previously validated this identical Rap1 HSV in multiple studies.^38,59^ HSV transgene expression in the brain peaks within a few days following infusion, with elevated levels lasting until about 6 days post-infusion and ceasing around 8 days post-infusion.^64,65^ This rapid rate of HSV-mediated transgene induction is well-suited to mimic the time period of our escalating, intermittent stress protocol. HSV-Rap1-GFP or HSV-GFP (as a control) were infused into the CA3 region of mice, and animal were transcardially perfused 4 days later (**Figure 5A**). As expected, the area of viral infection extended slightly beyond the CA3 region and into the neighboring CA2 region; however, CA1 neurons were rarely infected (**Figure 5B**). Identical to the previously described stress experiments, dendritic spine analysis was performed on CA3 pyramidal neuron secondary dendrites of the apical tree, and the n for statistical analysis is the number of mice assessed in each viral condition. 4-day Rap1 overexpression reduced the total density of dendritic spines in CA3 neurons [t(6)=2.830, p=0.0300] (**Figures 5C and 5D**). Subtype analysis revealed that Rap1 specifically reduced the density of thin spines [t(6)=3.186, p=0.0189] (**Figure 5E**), with no corresponding effects on either stubby [t(6)=0.08964, p=0.9315] (**Figure 5F**) or mushroom spines [t(6)=0.3280, p=0.7540] (**Figure 5G**). Further, survival curve analysis of head diameters within different spine subtypes indicates that Rap1 had no bearing on the head diameter distribution of thin (χ2=2.176, p=0.1401) (**Figure 5H**) or stubby spines (χ2=1.964 p=0.1610) (**Figure 5I**). On the other hand, Rap1 caused a slight, yet significant, rightward shift in the head diameter distribution curve of mushroom spines (χ2=6.357, p=0.0117) (**Figure 5J**). This rightward shift, in the absence of changes in the density of mushroom spines, suggests that Rap1 is increasing, albeit slightly, the head size of a preexisting population of mushroom spines.

**Figure 5.**
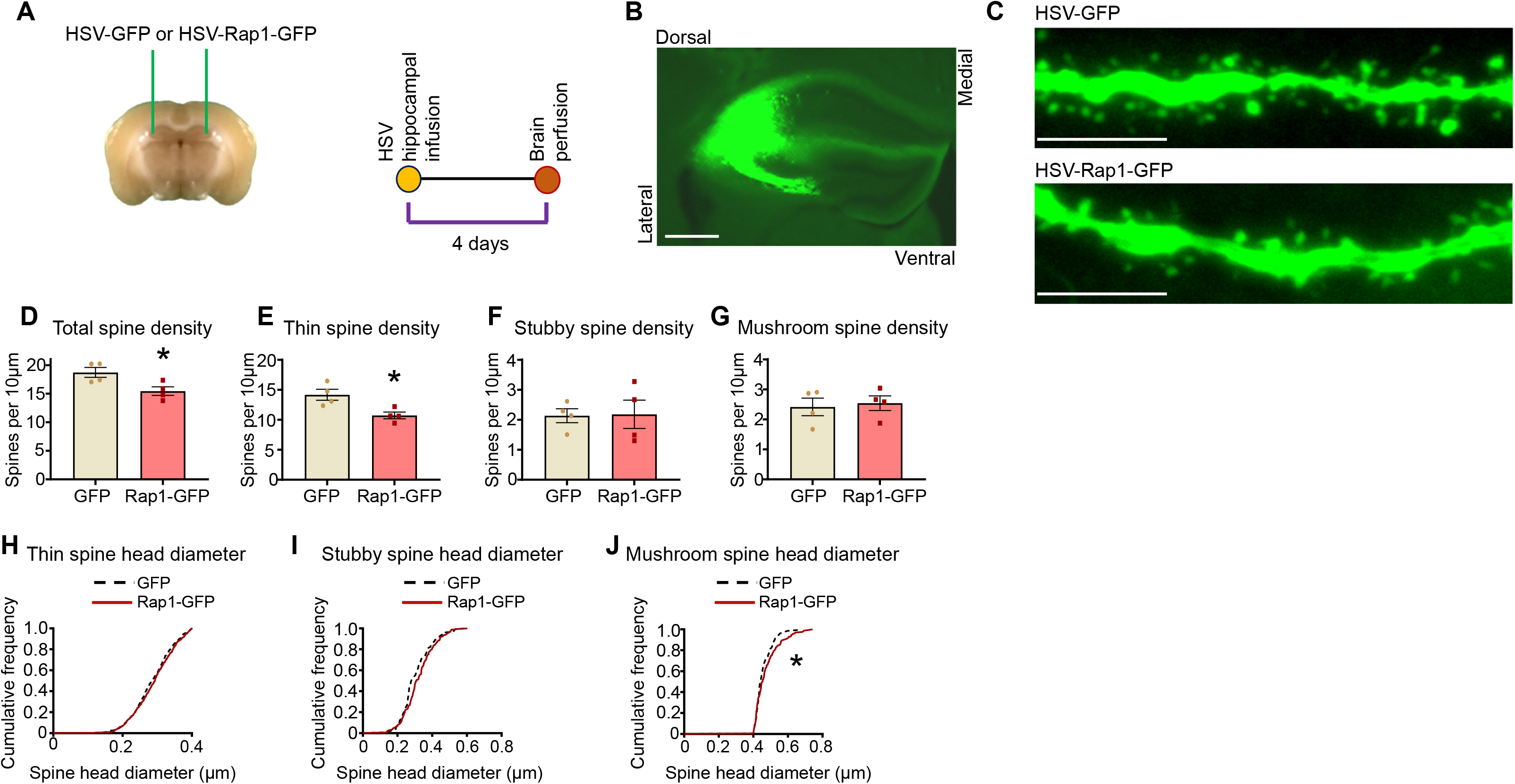
Rap1 reduces the stability of select dendritic spine subtypes in CA3 pyramidal neurons. A. Schematic of experimental design. Mice were infused with HSV co-expressing Rap1 and GFP (HSV-Rap1-GFP) or HSV-GFP into the CA3. 4 days post-viral infusion, mice were transcardially perfused. B. Representative low magnification image showing the left hemisphere hippocampus. Surgical coordinates used result in the targeting of HSV-GFP to the CA3 and CA2 region of the dorsal hippocampus. Scale bar=500µm C. Representative images of CA3 region pyramidal neuron secondary dendrites of the apical tree expressing GFP or Rap1-GFP. Scale bar=10µm. D. Graph depicts the total dendritic spine density of CA3 pyramidal neurons expressing GFP and Rap1-GFP. *Two-tailed t-test, p<0.05. 3-5 neurons were imaged per mouse and spine values across all neurons within a single mouse were averaged. n=4 GFP and 4 Rap1-GFP mice. E. Graph depicts thin spine density in CA3 pyramidal neurons expressing GFP or Rap1-GFP. *Two-tailed t-test, p<0.05. n is same as in 5D. F. Graph depicts stubby spine density in CA3 pyramidal neurons expressing GFP and Rap1-GFP. No significant differences were detected. n is same as in 5D. G. Graph depicts mushroom spine density in CA3 pyramidal neurons expressing GFP and Rap1-GFP. No significant differences were detected. n is same as in 5D. H. No significant differences in the cumulative thin spine head diameter curve were identified between the GFP and Rap1-GFP conditions. n=782 spines in HSV-GFP mice and 693 spines in HSV-Rap1-GFP mice. I. No significant differences in the cumulative stubby spine head diameter curve were identified between the GFP and Rap1-GFP conditions. n=124 spines in HSV-GFP mice and 146 spines in HSV-Rap1-GFP mice. J. A modest, yet significant rightward shift in the cumulative mushroom spine head diameter curve was identified in stressed mice vs. non-stressed mice. *Mantel-Cox, p<0.05. n=140 spines in non-stressed mice and 162 spines in stressed mice. All summary data are the mean + SEM.

### Key differences in the spatial distribution of overexpressed Rap1 in different dendritic spine subtypes

We found that escalating, intermittent stress increases Rap1 in CA3 synaptosomes and decreases the stability of thin dendritic spines. Further, the *in vivo* overexpression of Rap1 in CA3 neurons pheno-copies the effects of stress on reduced thin dendritic spine density. Although Rap1 is enriched in dendritic spines,^63^ owing to the limited imaging resolution of prior Rap1 synaptic studies, the precise distribution of Rap1 within dendritic spines is not known. As an initial step in understanding the selective vulnerability of thin spines in response to Rap1 overexpression, we examined the spatial distribution of overexpressed Rap1 in the dendritic spines of hippocampal neurons to determine if this distribution differs as a function of spine subtype. To this end, we used structured illumination microscopy (SIM) for the visualization of immunolabeled proteins within small domains of dendritic spine heads and necks, referred to as nanodomains.^66^ As SIM-mediated spatial mapping of dendritic spine proteins is optimal within culture systems and is far less amenable within *ex vivo* systems, we overexpressed GFP alone, or in combination with Myc-tagged Rap1 (Rap1b isoform), in mature cultured hippocampal neurons (DIV23). 2.5 days post-transfection, neurons were fixed and Myc immunolabeling was used to exclusively visualize overexpressed pools of Rap1, with attention devoted to dendritic spines located on secondary dendrites of the apical tree. We used 2.5 day overexpression to allow sufficient time to detect overexpressed pools of Rap1 while simultaneously limiting effects on baseline thin spine density that occur from longer term Rap1 overexpression.

We found that overexpressed Rap1 is not homogenously distributed throughout the spine, but rather is contained within discrete nanodomains throughout the neck and head of individual spines (**Figures 6A and S5A**). Thin spines, which based on our classification, have a smaller spine head area than mushroom spines (p<0.0001) (**Figure S5B)**, also have a smaller neck area than mushroom spines (p=0.0002) (**Figure S5C**), as well as a smaller total spine area than mushroom spines and filopodia (p<0.0001) (**Figure S5D)**. Quantification revealed that Myc-Rap1 is trafficked to about 75% of postsynaptic protrusions (i.e., spines and filopodia). To determine if protrusion gross morphology influences the likelihood that an individual protrusion contains pools of overexpressed Rap1, we examined the percentage of filopodia, thin, stubby, and mushroom spines that contain Myc-Rap1 nanodomains. We found that Myc-Rap1 nanodomains are rarely present in stubby spines (9%). In contrast, Myc-Rap1 nanodomains were present in the majority of filopodia (82%), thin spines (60%), and mushroom spines (89%) (**Figure 6B**). Within thin and mushroom spines, we noted that overexpressed Rap1 is trafficked to both the spine head and neck regions. To assess potential differences in the overall spatial distribution of Rap1 between thin and mushroom spines, we examined the percentage of each spine subtype that contain Myc-Rap1 nanodomains in the spine head, spine neck, or in both the head and neck. Overall, this analysis revealed that mushroom spine heads more often contain Myc-Rap1 nanodomains compared to thin spine heads (82% for mushroom, 47% for thin), while thin spine necks are more likely to contain Myc-Rap1 nanodomains than mushroom spine necks (54% for mushroom, 74% for thin) (**Figures 6C and 6D**).

**Figure 6.**
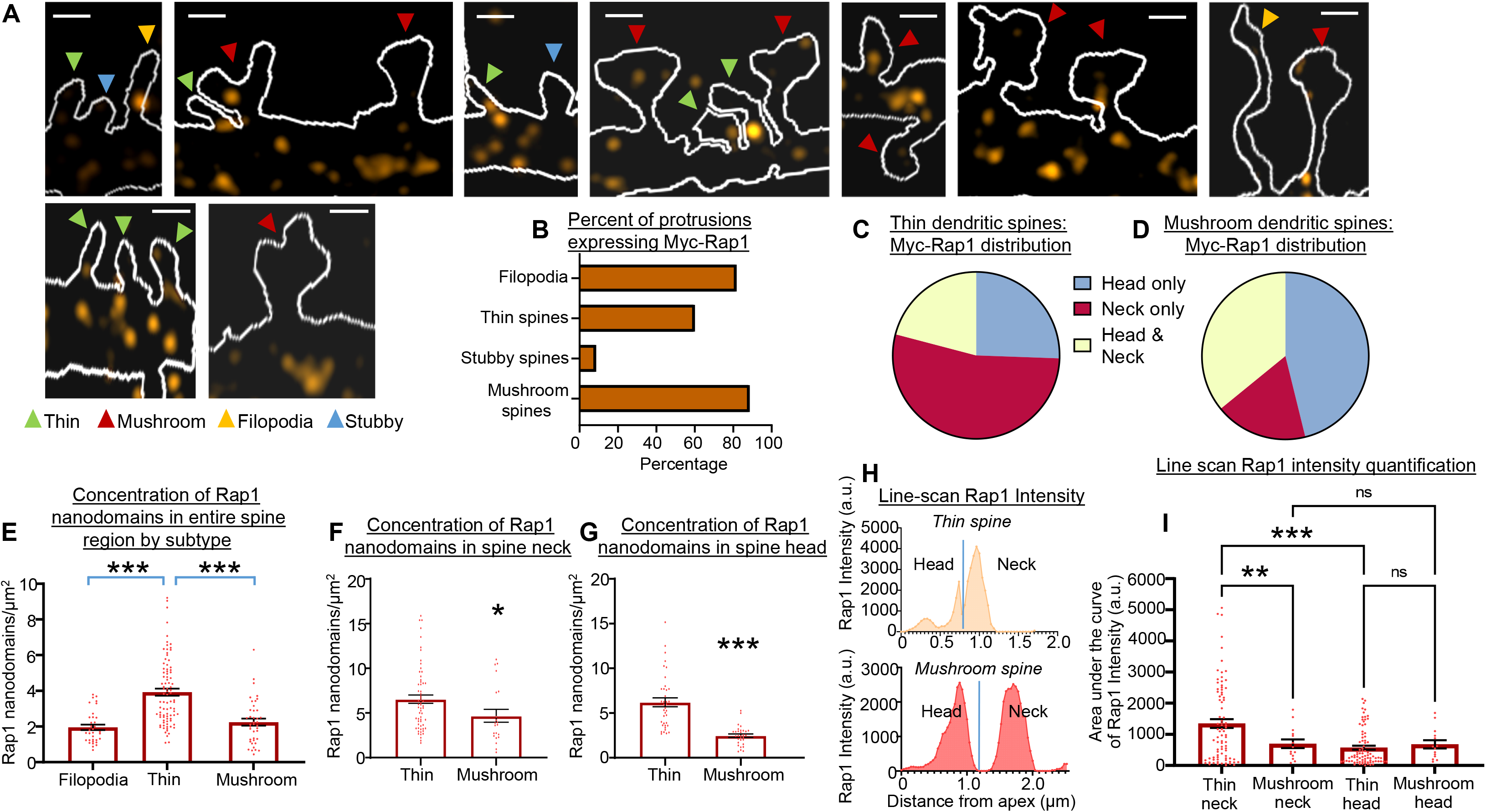
The spatial distribution of overexpressed Rap1 varies within different spine subtypes. A. Cultured hippocampal neuron SIM images of individual dendritic spines showing Rap1 nanodomains (Myc-tag immuno-labeled) in orange. Scale bar=1µm. B. Graph depicts the percent of filopodia and indicated spine subtypes that contain Myc-Rap1 nanodomains upon overexpression in mature cultured hippocampal neurons. C. Among thin spines containing overexpressed Rap1, the chart depicts the proportion located in each indicated spine region. D. Among mushroom spines containing overexpressed Rap1, the chart depicts the proportion located in each indicated spine region. E. Among the indicated protrusions exhibiting overexpressed Rap1, the graph depicts the nanodomain density of Myc-Rap1 throughout the entire filopodia or spine region (head and neck combined). ***Bonferroni, p<0.001. n=34 filopodia, 88 thin spines, 40 mushroom spines from 7 neurons. F. Among thin and mushroom spines exhibiting overexpressed Rap1 in the neck, graph depicts the nanodomain density of Myc-Rap1 in the neck. *Two-tailed t-test, p<0.05. n=61 thin spines and 21 mushroom spines from 7 neurons. G. Among thin and mushroom spines exhibiting overexpressed Rap1 in the head, graph depicts the nanodomain density of Myc-Rap1 in the head. ***Two-tailed t-test, p<0.0001. n=38 thin spines and 31 mushroom spines from 7 neurons. H. Representative line scans through a thin spine and a mushroom spine. Graph shows the intensity of the Rap1 signal as a function of distance across the head region and neck region for each spine subtype. I. Graph depicts the area under the curve of Rap1 intensity. Quantification is from a randomly selected population of thin and mushroom spines. One-way ANOVA with Dunnett’s multiple comparison correction, **p<0.01, ***p<0.001. n=90 thin neck, 14 mushroom neck, 90 thin head, 14 mushroom head; from 4 neurons. All summary data are the mean + SEM.

As thin spines have a significantly lower total surface area than filopodia and mushroom spines, we theorized that this lower area may restrict the number of Myc-Rap1 nanodomains contained within thin spines. Indeed, as expected based on their lower surface area, we found that thin spines have significantly fewer Myc-Rap1 nanodomains than either filopodia (Bonferroni p<0.0001) or mushroom spines (Bonferroni p<0.0001) (**Figure S6A**). We then used linear regression to determine if the total protrusion area within the individual subclasses of filopodia, thin spines, and mushroom spines is related to the number of nanodomains. Interestingly, we found that as the size of filopodia and thin spines increase, there is a corresponding increase in the number of Myc-Rap1 nanodomains present (correlation analysis, p=0.0034 for filopodia; p<0.0001 for thin spines) (**Figure S6B and S6C**). On the other hand, the size of individual mushroom spines had no significant bearing on the number of Myc-Rap1 nanodomains present (correlation analysis, p=0.23) (**Figure S6D**). Overall, this suggests that within thin spines and filopodia, but not mushroom spines, the protrusion size at least partially influences the amount of Myc-Rap1 present.

Next, we reasoned that although the prior analysis indicates that thin spines have fewer total Myc-Rap1 nanodomains than mushroom spines, the selective vulnerability of thin spines to the destabilizing effects of Rap1 could be related to differences in the concentration or density of Myc-Rap1 in thin spines compared to mushroom spines. More specifically, we theorized that Myc-Rap1 is potentially more concentrated in thin spines compared to mushroom spines. If so, this could indicate that the concentration of overexpressed Rap1 in mushroom spines is not sufficient to impact the stability of this spine class, while the comparatively higher concentration of Rap1 in thin spines is sufficient to attenuate stability. To test this, within the population of protrusions containing at least one Myc-Rap1 domain, we determine the proportion of the area of each protrusion subtype that is occupied by Myc-Rap1. Interestingly, we found that thin spines exhibit a greater proportion of surface area occupied by Myc-Rap1 than either mushroom spines (Bonferroni p=0.0005) or filopodia (Bonferroni p=0.0002) (**Figure S6E**). Linear regression within each protrusion subtype indicates that for filopodia and thin spines, as protrusion area decreases there is a corresponding increase in the proportion of the spine containing Myc-Rap1 (correlation analysis, p=0.045 for filopodia; p=0.0008 for thin spines) (**Figures S6F and S6G**). No corresponding relationship was identified for mushroom spines, however (correlation analysis, p=0.38) (**Figure S6H**).

Next, we wanted to determine if the increased concentration of Myc-Rap1 in thin spines as compared to mushroom spines is due to selective enrichment in the spine head and/or spine neck. To this end, we quantified the number of Myc-Rap1 nanodomains present per µm^2^ total spine area and per µm^2^ area of the spine neck and spine head. Among protrusions containing at least one Myc-Rap1 nanodomain, we found that thin spines contain a denser concentration of Myc-Rap1 throughout the entire spine as compared to filopodia and mushroom spines [one-way ANOVA (27.57), p<0.0001; Bonferroni thin vs. filopodia, p<0.0001; mushroom vs. thin, p<0.0001; filopodia vs. mushroom, p>0.999] (**Figure 6E**). As filopodia lack a discrete head region, assessments of Myc-Rap1 nanodomain concentration in head and neck regions were performed only for thin and mushroom spines. Among spines containing at least one Myc-Rap1 nanodomain in the neck region, we found that the nanodomain concentration was greater in the neck for thin spines compared to mushroom spine [t(80)=2.068, p=0.0419] (**Figure 6F**). Similarly, among spines containing at least one Myc-Rap1 nanodomain in the head region, we found that the nanodomain concentration was greater for thin vs. mushroom spines [t(67)=6.499, p<0.0001] (**Figure 6G**).

Although thin spines contain a greater density of Myc-Rap1 nanodomains in the head and neck region as compared to mushroom spines, this does not necessarily indicate that the integrated concentration of Myc-Rap1 is greater. Indeed, factors such as nanodomain size and the label intensity of individual nanodomains are also important factors. To quantify integrated levels of Myc-Rap, we performed line-scans through the head and neck of individual spines (**Figure 6H and Figure S7**). This analysis revealed that the area under the curve of Rap1 intensity within thin spines is greater in the neck region compared to the head region (one-way ANOVA, Dunnett’s p<0.0001), and that this Rap1 intensity is greater in thin spines necks compared to mushroom spine necks (one-way ANOVA, Dunnett’s p=0.0077) (**Figure 6I**). In contrast, the area under the curve of Rap1 intensity did not differ between thin spine heads and mushroom spine heads (one-way ANOVA, Dunnett’s p=0.9213 (**Figure 6I**). That our analysis identified an increased nanodomain concentration in thin spine heads vs. mushroom spine heads, yet the area under the curve of Rap1 intensity in the head region of these spine subtypes does not differ, could arise due to nanodomains in thin spine heads being of a smaller average size and/or lower average intensity. Collectively, these findings pinpoint the heightened integrative concentration of Myc-Rap1 in the neck of thin spines as a major distinguishing feature relative to that of mushroom spines.

### Rap1 overexpression does not affect hippocampal neuron dendrite parameters

In addition to spines, our SIM analysis of cultured hippocampal neurons indicates that following overexpression, Myc-Rap1 nanodomains are also highly prevalent within dendrite shafts (**Figure S8A**). We were thus interested in determining if Rap1 overexpression impacts the arborization/branching of apical and basal dendrites in cultured hippocampal neurons. Surprisingly, we found that Myc-Rap1 overexpression in mature cultured hippocampal neurons did not impact the length of the apical or basal dendrite trees (**Figure S8B-S8D**) and similarly did not impact the number of terminal apical or basal dendrites (**Figure S8E and S8F)**. Further, Sholl analysis of the basal and apical dendrite trees did not detect any impact of Rap1 overexpression on dendrite complexity (**Figure S8G and S8H**).

### Rap1 overexpression in the CA3 region affects object-in-place associative recognition memory

Using the identical escalating, intermittent stress protocol detailed here, we previously showed that stress impairs object-in-place memory.^38^ Object-in-place memory is a form of associative recognition memory, as it requires mice to form a lasting association between objects and their relative locations, assessed by the recognition of a change in the location of previously encountered objects after a delay period.^67^ Object-place memory associations involve processing occurring in the CA3 and are highly sensitive to manipulations of this region.^68,69^ To determine if increased Rap1 in the CA3 is sufficient to affect object-in-place memory, we infused either HSV-GFP or HSV-Rap1-GFP into the CA3 of mice. We then assessed the ability of mice to recognize a change in the location of two objects, which manifests as a preference to investigate objects that swapped locations between trials vs. objects whose location was fixed between trials (**Figure 7A**).

**Figure 7.**
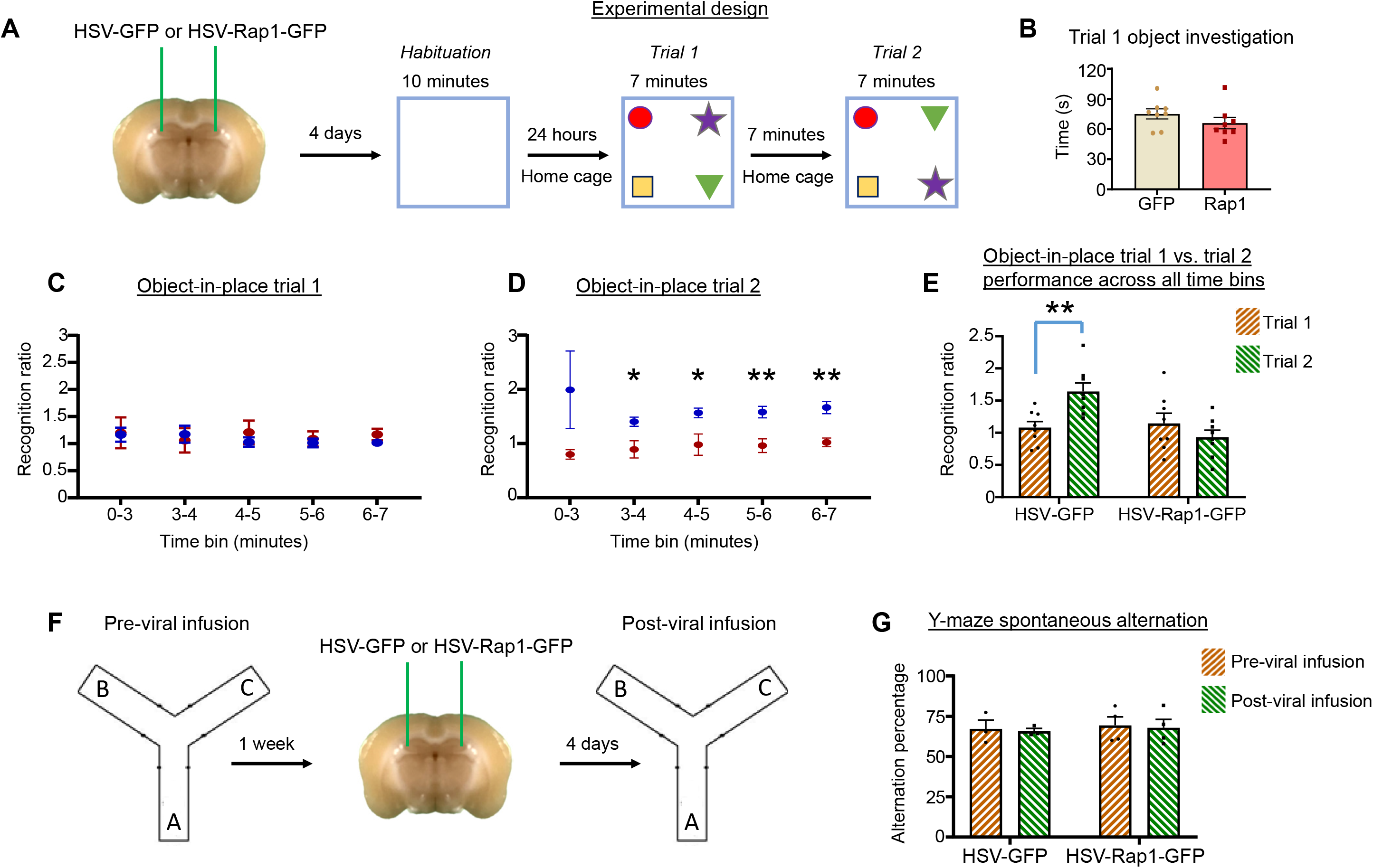
Overexpression of Rap1 in the CA3 region impairs recognition memory. A. Schematic of experimental design. Mice were infused with HSV-Rap1-GFP or HSV-GFP into the CA3. 4 days later mice underwent habituation in an open field. The next day mice were assessed for object-in-place recognition memory. During trial 2, mice should prefer to explore the objects that swapped locations compared to trial 1 (purple star and green triangle in the schematic). B. Graph depicts time GFP and Rap1-GFP mice spent exploring objects during trial 1 of the object-in-place task. No differences between groups were identified. n=8 GFP and 8 Rap-GFP mice. C. Graph depicts the trial 1 object-in-place recognition ratio for the GFP and Rap1-GFP mice. No differences between groups were identified at any time bins. n=8 GFP and 8 Rap-GFP mice. D. Graph depicts the trial 2 object-in-place recognition ratio for the GFP and Rap1-GFP mice. Differences were identified at all time bins except the 0-3 minute bin. *p<0.05, **p<0.01. n=8 GFP and 8 Rap-GFP mice. E. Graph depicts recognition ratio across all 7 minutes of trial 1 and trial 2 of the object-in-place task. **p<0.01. n=8 GFP and 8 Rap-GFP mice. F Schematic of experimental design for Y-maze testing. Mice were assessed for spontaneous alternation in the Y-maze 1 week prior to HSV infusion and 4 days post-HSV infusion. G. HSV-Rap1 did not affect spontaneous alternation performance post-HSV infusion relative to the pretest. n=3 GFP, 4 Rap1 mice. All summary data are the mean + SEM.

Accurate assessments of object-in-place memory require that mice have a normal drive to explore novel objects. Thus, we first assessed the amount of time mice spent investigating objects throughout trial 1 as an index of novel exploration motivation. We found that the GFP and Rap1-GFP groups spent nearly identical amounts of time exploring objects during trial 1 [t(14)=1.199, p=0.2503], indicating that Rap1 in the CA3 does not impact motivation for novelty exploration (**Figure 7B**). During the second trial, to assess the ability of mice to recognize the change in object locations, we calculated a recognition ratio with chance performance equal to 1.0 (see methods). For trial 2, the cumulative recognition ratio was assessed for the first three minutes and for each minute thereafter. This same calculation was used for trial 1 to ensure that mice did not have any pre-existing preferences for either pair of objects. As expected, during trial 1 there were no differences in the recognition ratio between the GFP and Rap1-GFP groups across the 7 minute period [Rap1 main effect, F(1,14)=0.1261, p=0.7278], or within any time bins (post hoc p values: 0-3min p=0.9609, 3-4min p=0.9609, 4-5min p=0.9161, 5-6min p=0.9609, 6-7min p=0.7716) (**Figure 7C**). Further, during trial 1 neither group’s performance differed from a chance level of 1.0 (GFP p=0.4177, Rap1 p=0.3851), indicating that the mice did not have any preexisting preferences for any object pairs. On the other hand, for trial 2 we found that global performance across the 7 minutes was impaired in the Rap1-GFP group compared to the GFP group [Rap1 main effect, F(1,14)=17.23, p=0.0010]. Further, whereas the GFP group performed above chance levels (p=0.0018), the Rap1-GFP group did not (p=0.3851) (**Figure 7D**). Post hoc testing of individual time bins in trial 2 revealed a decrease in the recognition ratio in the Rap1-GFP group as compared to the GFP group at all time bins except the 0-3 minute bin (post hoc p values: 0-3min p=0.1417, 3-4min p=0.0478, 4-5min p=0.0478, 5-6min p=0.0089, 6-7min p=0.0025) (**Figure 7D**). Additionally, we assessed the raw times individual mice spent investigating the swapped and non-swapped object pairs during trial 2, which confirmed the impaired performance in the Rap1 group (**Figures S9A and S9B**).

As the prior statistical analyses assessed performance in trial 1 and trial 2 separately, we also performed a repeated measures analysis of recognition ratios across the 7 minute period for trial 1 and trial 2. We found a main effect for Rap1 overexpression status [F(1,14)=8.489, p=0.0113] and a significant interaction between Rap1 overexpression and trial [F(1,14)=7.881, p=0.014]. Post hoc analysis indicated that the GFP group showed an increased preference for the swapped object pair during trial 2 compared to the investigation of these same objects during trial 1 (Bonferroni post hoc, p=0.0008), whereas the Rap1-GFP group showed no change in preference (Bonferroni post hoc, p=0.7168) (**Figure 7E**). Taken together, these findings indicate that Rap1 overexpression in the CA3 causes a striking impairment in object-in-place associative recognition memory.

### Rap1 overexpression in the CA3 regions does not affect spontaneous alternation behavior

In addition to object-in-place deficits, our prior findings indicate that the identical escalating, intermittent stress procedure used in the present study also impairs spontaneous alternation performance in the Y-maze.^38^ Y-maze spontaneous alternation is a commonly used measure of spatial working memory^70^ and is CA3 dependent in mice.^71^ To assess the impact of Rap1 overexpression in the CA3 on spontaneous alternation, we used a repeated measures design in which mice were assessed in the Y-maze one week prior to HSV infusion (trial 1) and re-assessed 4 days post-HSV infusion into the CA3 (trial 2) (**Figure 7F**). In both trials, mice were allowed 10 minutes of continual exploration of the three maze arms. The spontaneous alternation percentage was calculated for both trials; chance performance is equal to 50% (see methods). Comparing performance in trial 2 (viral transgene expression in the CA3 onboard) vs. trial 1 (prior to viral infusion), we found that Rap1 overexpression in the CA3 has no impact on spontaneous alternation performance (2-way ANOVA with Bonferroni post hoc, Rap1 p>0.9999) (**Figure 7G**).

## Discussion

To our knowledge, these findings represent the first investigation into the effects of escalating, intermittent stress on hippocampal dendritic spines and associated biochemical alterations. Taken together, these findings illuminate a potentially key role for Rap1 in stress-mediated hippocampal synaptic aberrations as well as in specific aspects of stress-induced cognitive impairment. The potential clinical significance of these findings is extended by previous studies that have identified variants and mutations in genes encoding direct activators and inhibitors of Rap1, as well as altered forebrain expression of Rap1 itself (including in the hippocampus), in disorders known to be in many cases precipitated by stress, including anxiety disorders, major depressive disorder, and schizophrenia.^72–77^

Similar to the present study in hippocampal neurons, our prior study demonstrated that escalating, intermittent stress reduces dendritic spine density in mPFC neurons.^38^ However, there are key differences in the predominant subtype affected in each brain region. Notably, escalating, intermitted stress reduces the density of mushroom spines in mPFC neurons, with no effect on thin or stubby spines,^38^ while it reduces the density of thin spines in CA3 neurons, with no effects on other subtypes. Importantly, the same procedures were used in both studies to illuminate and quantify spines, thus, subtype differences are not likely due to methodological influences. Comparison of these studies identified a notably greater baseline density of thin spines in CA3 vs. mPFC pyramidal neurons, with no apparent differences in the density of stubby or mushroom spines. Whether this increase in baseline thin spine density in CA3 neurons increases the influence of stress on this spine subtype relative to mPFC neurons is not clear. Consistent with escalating, intermittent stress causing a reduction in spine density within CA3 neurons, we found this stress procedure also reduces levels of surface GluA1 receptors in the CA3 region. Whether stress-mediated dendritic spine reductions in the mPFC are similarly mirrored by a regional reduction in surface GluA1 content is currently not known, but remains an interesting question for the future.

Our prior study found that escalating, intermittent stress increases Rap1 in mPFC synaptosomes,^38^ and here we find a similar effect in CA3 synaptosomes, with no differences in other examined small GTPases. Interestingly, Rap1 overexpression has differing spine subtype effects in pyramidal neurons of the CA3 vs. mPFC. More specifically, although Rap1 overexpression in both regions promotes spine destabilization as evidenced by a significant reduction in total dendritic spine density, this reduced total density is due to the selective reduction of thin spines in CA3 pyramidal neurons and due to selective reductions in mushroom spines in mPFC pyramidal neurons.^38^ Of importance is that the spine effects of Rap1 overexpression in each region mirror the effects of escalating, intermittent stress on spine subtype destabilization in each region. Indeed, escalating intermittent stress increases Rap1 in CA3 synapses and causes a reduction in thin spines, and viral-mediated overexpression of Rap1 in the CA3 also causes a selective reduction in thin spines. On the other hand, escalating, intermittent stress increases Rap1 in mPFC synapses and causes a reduction in mushroom spines, and viral-mediated Rap1 overexpression causes a selective reduction in mushroom spines in the mPFC.^38^ The reasons for the divergent effects of Rap1 overexpression in the CA3 vs. mPFC on spine subtype stability remain unclear. It is possible that the trafficking of Rap1 into different types of spines varies in mPFC neurons relative to those of the CA3.

Interestingly, we found that upon overexpression in hippocampal pyramidal neurons, Rap1 is not uniformly trafficked into all spine subtypes. Rather, stubby spines rarely accumulate overexpressed Rap1, while the majority of thin and mushroom spines contain overexpressed Rap1. When comparing thin spines and mushroom spines, thin spines show a greater proportion of their surface area occupied by Rap1 than mushroom spines, and thin spines also show a greater concentration/density of Rap1 nanodomains than mushroom spines. Integrated Rap1 intensity assessments indicate that within thin spines, overexpressed Rap1 is more concentrated in the neck region compared to the head region, and that the major difference in the concentration of Rap1 between thin spines and mushroom spines lies in the neck region. More specifically, thin spine necks contain a greater concentration of integrated Rap1 signal than those of mushroom spines, while the integrated Rap1 signal in the head region was similar between thin and mushroom spines. The neck of spines is critical for both biochemical and electrical compartmentalization, and spine neck morphology has major influences on broader neuronal excitability.^78,79^ Similar to the head, the neck of spines is enriched in filamentous actin (F-actin). In the neck, this F-actin is mainly arranged longitudinally, and is comprised of branched and linear actin.^80^ Rap1 is capable of severing F-actin through its influence on the activity of the actin binding protein cofilin.^81,82^ It is conceivable that upon overexpression the particularly elevated concentration of Rap1 in the neck of thin spines leads to regional actin destabilization, thereby increasing the likelihood of neck collapse and therefore reduced thin spine numbers.

It is possible that Rap1 overexpression fails to alter the density of mushroom spines due to the concentration of overexpressed Rap1 in mushroom spine necks and/or heads being insufficient to impact the stability of this spine class. It is also possible that the protein activity of Rap1 differs when present in mushroom spines vs. thin spines. Rap1 directly binds to several PDZ domain-containing scaffolding proteins, including SHANK3 and AF6 (afadin), both of which have an established enrichment in dendritic spines.^41–43^ Evidence suggests that AF6 affects the trafficking of Rap1 into dendritic spines, and impacts Rap1’s ability to affect spine morphogenesis.^43^ Whether differential interactions with AF6 contribute to the observed differences in the anchoring of overexpressed Rap1 in different spine subtypes and within different spine domains (e.g., head and neck) remains an interesting question for future endeavors.

It is possible that the selective cognitive impairments resulting from Rap1 overexpression in the CA3 are a product of the specific effects of Rap1 in destabilizing thin and not mushroom spines. Mushroom spines typically contain more functional AMPA receptors than thin spines.^53,83^ Owing to their small head size and low content of functional AMPA receptors, thin spines exhibit a greater potential for plasticity in response to synaptic engagement than mushroom spines. Indeed, when stimulated, thin spines can gradually transition to mushroom spines, while stimulation of mushrooms spines typically results in only a very short-lived increase in head size, with eventual reversion back to the pre-existing size.^84^ Based on these considerations, it is theorized that thin spines are “learning spines” due to their unusually high malleability in response to neural inputs, while mushroom spines are “memory spines” due to their persistence over time thereby allowing for a stable synaptic representation of learned experiences.^85^ Spine head enlargement occurring within thin spines in response to synaptic engagement is very rapid, occurring within minutes,^84^ and this rapid molding is likely of critical importance for the involvement of thin spines in new learning.

Our escalating, intermittent stress protocol impairs object-in-place associative recognition memory and Y-maze spontaneous alternation.^38^ Although the CA3 region is involved in both object-in-place associative memory and spontaneous alternation,^68,69,71^ Rap1 overexpression in CA3 neurons only affected object-in-place memory. The object-in-place task requires the learning of object locations over a 7 minute exploration period and the subsequent assessment of this memory after a 7 minute delay. It is conceivable that the learning of the object locations is due, at least in part, to the engagement of thin “learning spines,” and thus manipulations that reduce the baseline density of thin spines could impede the learning capabilities needed for this task. Y-maze spontaneous alternation requires mice to maintain and constantly update a mental log of arm entry choices. This process is continually occurring throughout the 10 minute fee choice period. In mice, the timeframe for working memory is very brief as the information held in mind is only useful for a short duration. Indeed, Y-maze arm entry choices made several minutes prior will have no useful bearing on a mouse’s upcoming arm entry selections. In this way, it is conceivable that spontaneous alternation is less dependent on “learning spines” as this task does not require longer term or even multi-minute maintenance of specific arm entry selection choices.

We found that whereas control mice exhibit an increase in c-Fos integrated intensity in the CA3 following object exploration, mice subjected to escalating, intermittent stress did not show a corresponding increase in c-Fos intensity. This suggests that stress induces a state of reduced CA3 responsivity or engagement in response to novelty exploration. This reduced engagement during the initial phases of object exploration could hinder the ability of mice to detect changes in the location of objects during subsequent exposures, and thus could contribute to our previously observed effects of escalating, intermittent stress on impaired object-in-place performance.^38^ Whether the effects of stress on diminished c-Fos integrated intensity in the CA3 is due in part to our identified selective reduction in thin spines along CA3 pyramidal neurons is not known. Further, our prior work did not determine if escalating, intermittent stress, which in the mPFC selectively reduces mushrooms spines, attenuates the effects of novelty exploration on c-Fos levels in the mPFC. Knowledge of this would help determine if mushroom spines reductions are also associated with reduced regional responsivity to novelty exploration. If not, based on our current findings, this would be suggestive of a level of specificity for thin spine reductions in potentially contributing to regional responsivity aberrations.

Our prior study^38^ together with the present findings implicate heightened synaptic levels of Rap1 in contributing to the effects of stress on mPFC and hippocampal neuronal structural and cognitive aberrations. Although we found no changes in the expression of other examined small GTPases in either brain region, other studies have implicated additional small GTPases as potentially key mediators of the neural response to stress. Notably, social defeat stress decreases dendritic spine density in mPFC neurons and causes an increase in the enzymatic activity of a particular Rap isoform, Rap2.^86^ 10 days of consecutive restraint stress decreased the enzymatic activity of Rac1 in hippocampal whole cell lysates and increased Rac1 activity in basolateral amygdala whole cell lysates; the activity of RhoA was not altered in either brain region.^87^ 14 days of consecutive stress increased total levels of RhoA protein in the hippocampus 1 day post-stress with no effect on Rac1/2/3 levels.^88^ A single stress exposure, on the other hand, had no effect on RhoA protein in the hippocampus.^88^

Although we found that the stress increases Rap1 levels in the CA3 and that overexpression of Rap1 in the CA3 region is sufficient to cause stress-relevant phenotypes, including cognitive deficits, a limitation is that we did not determine if the knockdown of Rap1 in the CA3 prevents the effects of stress on the various neural phenotypes observed. This was not assessed as the knockdown or interference of Rap1 within the forebrain, including within individual forebrain regions, produces cognitive impairments, including impaired spatial memory, impaired context discrimination, impaired memory retrieval, and impaired object-in-place memory.^38,89,90^ The ability to determine if Rap1 is essential for stress-mediated cognitive impairment would require that baseline cognition is relatively normal when Rap1 levels are reduced, which is not the case. Similarly, reducing Rap1 activity causes a striking alteration in pyramidal neuronal dendritic spine morphology such that spine area more than doubles with most spines exhibiting a mature, mushroom-like appearance.^43^ This baseline shifting of spine morphology to a mature state would likely preclude reliable assessments of the ability of stress to increase the density of thin spines in neurons with reduced Rap1, as stress would have to overcome a separate factor pushing spines toward a mushroom phenotype and away from a thin phenotype.

Importantly, our findings revealed that the effects of escalating, intermittent stress on synaptosomal Rap1 levels in the CA3 are sex-specific, as stress increased Rap1 in male, but not female, mice. The sex-dependent biochemical effects are in keeping with a growing body of rodent studies indicating that the response to stress, including restraint stress, differs considerably between sexes, and that these differences also extend to the molecular and biochemical signatures of stress in the hippocampus. Notably, recent work revealed that multi-week restraint reduces preference for sucrose in male rats, suggestive of anhedonia, with no comparable effect in female rats. Further, this same study found that restraint stress reduces protein levels of TrkB and GluA1 in the dorsal hippocampus of male rats, with no effect in female rats.^91^ Some studies suggest that estrogen protects female rodents from exhibiting many of the behavioral and biochemical aberrations that accompany stress. For example, chronic restraint stress reduces recognition memory in male, but not female, rats, which is associated with a male-specific reduction in surface GluA1 levels in the prefrontal cortex. Stress-mediated cognitive and biochemical phenotypes in female rats can be revealed upon interference with estrogen receptors.^92^ However, recent work indicates that female mice are protected from the effects of stress on cognition and on hippocampal dendritic spine loss during periods of low physiological estradiol (estrus), yet are susceptible to these stress effects during periods of high physiological estradiol (early-proestrous).^93^ This suggests that estrogen is not unconditionally protective against the effects of stress, and the consequences of estrogen reduction on stress-mediated phenotypes might differ if this reduction is due to physiological fluctuations that accompany the estrous cycle vs. reductions due to non-physiological pharmacological-mediated blockage. Future investigations aimed at determining if Rap1 levels in hippocampal synaptosomes are influenced by physiological fluctuations in estrogen and/or by exogenously-induced alterations in estrogen signaling, could offer insight into the mechanisms responsible for the sex-dependent Rap1 effects we identified.

In summary, our findings advance a growing narrative for the altered forebrain regulation of small GTPases in response to varying types of stress. Owing to their potent ability to influence the actin cytoskeleton within dendritic spines,^39^ small GTPases are strategically positioned to mediate the effects of stress on synaptic destabilization. One limitation of many prior studies investigating the effects of stress on small GTPases is that the ability of identified small GTPase alterations to produce stress-relevant synaptic and/or behavioral impairments is seldom known. Here we not only identify altered synaptic levels of hippocampal Rap1 in response to stress, but also determine that this alteration in Rap1 is sufficient in and of itself to produce hippocampal spine reductions and cognitive impairment. Whether the altered expression and/or activity of other small GTPases identified in other studies mimic the synaptic and cognitive effects of stress remains an open question for future endeavors.

## Acknowledgements

This work was supported by the National Institute of Health (National Institute of Mental Health) award R21MH125227 (MEC) and UW-Madison ICTR Award (MEC).

## Author Contributions

Behavioral experiments were performed by KJB and MEC. Biochemistry experiments and analyses were performed by KJB, AMV, and BAK. SIM imaging was performed by KJB. Dendritic spine imaging and analysis were performed by KJB and AMV. Dendrite imaging and analysis were performed by KJB and YY. KJB and MEC wrote the paper.

## STAR Methods

### Resource availability

#### Lead contact

Further information and requests for resource and reagents should be directed to and will be fulfilled by the Lead Contact, Michael Cahill.

#### Materials availability

This study did not generate new unique reagents. The HSVs used have been described previously,^38,59^ but were re-propagated for this study.

#### Data and code availability

This study did not generate code, but datasets related to the current study are available from the Lead Contact, Michael Cahill, upon request.

### Experimental model and subjects details

#### Experimental mice

c57BL/6J mice were used for all experiments (breeders from Jackson Laboratories). Male mice were used for all experiments except for the female mouse biochemical analysis as indicated in the main text and in the corresponding supplementary figure. All mice were between 10-13 weeks of age at the time of experimentation, with age matching between experimental groups. Mice were housed in standard sized cages with 4-5 mice per cage, and mice had access to food and water ad libitum. Mice weighed between 23-30 grams at the start of experiments. All mice were housed in a rodent vivarium located in the same building as the performed experiments, and mice were on a 12 hour light/dark cycle and maintained in a temperature-controlled environment. Simple randomization was used to assign mice to experimental groups. All of the experiments using animals were approved by the University of Wisconsin at Madison Animal Care and Use Committee.

#### Dissociated cortical neuron cultures

Dissociated cultures of primary hippocampal neurons were prepared from embryonic day 18 Sprague-Dawley rat embryos. Due to the age at time of harvesting, the determination of the sex of the embryos was not possible. Neurons were transfected between 21-23 DIV and were 25-28 DIV at the time of fixation or collection.

### Method details

#### Restrain stress

For restraint stress mice were immobilized in a broome-style mouse restrainer purchased from Plas-Labs (Cat. #551-BSRR). Mice underwent 6 bouts of immobilization stress over an 8 day period. On the first day mice were restrained for 20 minutes, followed by 2 hours of restraint the following day. Mice then had two consecutive days without restraint. This was then followed by four consecutive days of 6 hour restraint. Mice were randomly assigned to the restraint stress condition or to the non-stress control condition.

#### Sodium dodecyl sulfate-polyacrylamide gel electrophoresis (SDS-PAGE) and western blotting

Protein assays were performed on the S2, P1, and P2 fractions (BCA assay), and within a given fraction equal amounts of total protein for all samples were loaded onto a 4-15% Tris-HCL precast gel (BioRad). Samples were resolved via SDS-PAGE at 130 volts for 1.5 hours. Proteins were then transferred onto PVDF membranes for 1 hour at 100 volts (4⁰C) in 15% methanol transfer buffer. PVDF membranes were then air dried for 1 hour to allow for proteins to fully settle, rehydrated in methanol and blocked in Tris Buffered Saline (TBS) containing 5% bovine serum albumin (BSA) for 1 hour. Primary antibodies were diluted in TBS containing 5% BSA with overnight incubation. Peroxidase conjugated secondary antibodies were diluted in the same blocking buffer as the primary antibody with incubation at room temperature for 1 hour. Membranes were incubated with ECL substrate for 5 minutes (chemiluminescence) and developed using autoradiography film and machine processed using an All-Pro Imaging Corporation developer (model 100 plus). Protein levels were determined using densitometry using Image J. Protein levels were normalized to the corresponding levels of actin.

As equal amounts of total protein within a given fraction were used across all experimental samples, and because we found that actin levels were very consistent across experimental conditions, this indicates that actin levels are not altered by any of the experimental conditions and thus it is an appropriate protein for normalization. The following primary antibodies were used: Actin (Fisher Scientific, Cat# PIMA515739; 1:5000 dilution), Rac1 (Cytokeleton, Cat# ARC03; 1:1000 dilution), Cdc42 (Cell Signaling, Cat# 2466: 1:1000 dilution), Rap1 (Cell Signaling, Cat# 2326; 1:1000 dilution), and Ras (detects K-Ras, H-Ras, and N-Ras; Cell Signaling, Cat# 3339; 1:1000 dilution).

#### Subcellular fractionation

The fractionation of brain tissue into the P1, S2, and P2 fractions was performed using our previously detailed methods.^38,59,94,95^ Brains were rapidly sectioned into 1 mm thick coronal sections using an ice-cold metal brain matrix. The CA3 was dissected in coronal sections using a 0.22 mm micropuncher. The CA3 tissue micropunches were homogenized at 800 rpm in a Teflon homogenizer (15 up and down strokes) in buffer comprised of 0.32M sucrose, 4mM HEPES, with the addition of protease and phosphatase inhibitors (HEPES-sucrose buff22er). Protease/phosphatase inhibitor cocktail was purchased from Thermo Scientific (Cat# PI78440). The CA3 homogenate was then centrifuged at 1,000 x g (4⁰C) for 10 minutes. The resulting supernatant is the S1 fraction, and the resulting pellet the unwashed P1 fraction pellet. The P1 pellet was resuspended in HEPES-sucrose buffer, re-centrifuged at 1,000 x g (4⁰C) for 10 minutes resulting in the formation of a washed P1 pellet. The washed P1 pellet was then again resuspended in HEPES-sucrose buffer and the centrifugation process repeated two more times in order to obtain a purified P1 fraction which was then resuspended a final time in HEPES-sucrose buffer then snap frozen and stored at -80⁰C. The S1 fraction was centrifuged at 1,000 x g (4⁰C) and the supernatant collected – this step was repeated two more times in order to make sure no residual P1 pellet fragments are present in the S1 fraction. The S1 fraction was then centrifuged at 10,000 x g (4⁰C) for 15 minutes to produce a S2 supernatant and an unwashed P2 pellet. The S2 supernatant was removed and snap frozen and stored at -80⁰C. The unwashed P2 pellet was resuspended in HEPES-sucrose buffer and re-centrifuged at 10,000 x g (4⁰C) for 15 minutes. The resulted in a washed P2 pellet, which was again resuspended in HEPES-sucrose buffer and centrifuged a final time at 10,000 x g (4⁰C) for 15 minutes. The resulting P2 pellet was the resuspended in HEPES-sucrose buffer, then snap frozen and stored at -80⁰C. Just prior to SDS page, sample buffer containing SDS and DTT was added to each thawed solubilized fraction and samples heated to 95⁰C for 5 minutes.

#### Stereotaxic viral infusions into the CA3 region

HSV-Rap1-GFP was previously described and validated by our group in prior studies.^38,59^ Rap1 is expressed from an IE 4/5 promoter and GFP from its own CMV promoter. HSV-GFP (GFP expression from CMV promoter) was used for baseline comparisons. Mice were anesthetized with ketamine/xylazine, and virus infused into the CA3 using Hamilton 33-guage syringe needles with an infusion rate of 0.1 µl per hemisphere per minute for 5 minutes thereby resulting in the infusion of 0.5 µl per hemisphere. The was followed by a 5 minute waiting period in which the syringe needle remained in place despite no infusion occurring in order to allow for virus diffusion. The following stereotaxic coordinates, relative to Bregma, were used to specifically target the CA3 region of the hippocampus: anterior/posterior, -1.9mm; medial/lateral, +2.3mm; dorsal/ventral, -2.2mm.

#### BS3 (bis(sulfosuccinimidyl)suberate) Crosslinking

BS3 (bis(sulfosuccinimidyl)suberate) Crosslinking was performed using methods adapted from previous work.^54^ 24-hours following restraint stress or a non-stress control condition, mouse brains were dissected and rapidly sectioned into 1 mm thick coronal sections using an ice-cold metal brain matrix. The CA3 was dissected using a 0.1 mm micropuncher The CA3 punch was added to a 1.5mL Eppendorf rube containing 1mL of artificial cerebral spinal fluid (aCSF) and 40µl of 52mM BS3 was immediately added before being placed on a rocker at 4°c for thirty minutes. The timing of all incubations was kept identical between samples. After thirty minutes, the BS3 reaction was quenched by adding 100μl of 1M Glycine stock (for a final concentration of 100mM) and rocked for 10 minutes at 4°c. Samples were centrifuged at 20,000 x g (at 4°C) for 2 min to recover tissue. The tissue was resuspended in 50ul of ice-cold buffer containing 0.32M sucrose, 4mM HEPES, and protease/ phosphatase inhibitors (Cat# PI78440) before being sonicated. Samples were centrifuged at 20,000 x g (at 4°C) for 2 minutes to remove the insoluble material and the supernatant was collected, snap frozen, and stored at -80⁰C.

#### Diolistic labeling of neurons

Brains were transcardially perfused with phosphate buffered saline (PBS) for 1 minute using a flow rate of 5 mls per minute. This was then followed by perfusion with 1.5% paraformaldehyde for 7 minutes at flow rate of 5 mls per minute. Brains were then post-fixed in 1.5% paraformaldehyde for 1 hour. Brains were then coronally sectioned (150 µm thick) at room temperature (Vibratome). A Helios Gene Gun was used to biolistically deliver tungsten particles coated with DiI (micro-carriers) to coronal sections containing the CA3 field. Micro-carriers were made by combining 1.3µm tungsten particles (Bio-Rad, Cat# 1652268) with DiI and subsequent drying using a steady nitrogen flow for 20 minutes into a tubing preparation station (Bio-Rad, Cat# 1652418). Following biolistic delivery of the micro-carriers to the CA3, sections were incubated for 24 hours in room temperature PBS to allow for DiI diffusion throughout neurons. This was followed by a 10 minute incubation in 4% paraformaldehyde to seal the DiI in neurons. Sections were then washed in PBS and hardset vectashield (Vector Labs) used for coverslipping.

#### Dendritic spine imaging and analysis

Biolistically DiI labeled coronal sections were imaged as is. Sections from mice infused with HSVs underwent immunohistochemistry to boost the native GFP signal from the HSV using a chicken anti-GFP antibody from Aves Labs (Cat# GFP-1010, 1:3,000 dilution). Dendritic spines were imaged using a Keyence BZ-X700E scanning microscope. Images of secondary dendrites from the apical tree were captured using a 100x objective, with a minimum 40µm dendrite segment per neuron. Z-stack images were collapsed and a full-focus algorithm used to deconvolve the spine images. The NeuronStudio program was used for all dendritic spine quantification using methods detailed in our previous work.^38,49,52^ The experimental manually clicks onto the heads of individual dendritic spines resulting in the program placing a hollow ellipse onto each spine head such that the diameter of the ellipse corresponds to the diameter of the spine head. Manual adjustments can be made to the program-determined ellipse size in instances in which the ellipse under or over expands the true area of the spine head. The total number of spines and the head measurements of individual spines are then logged into a spreadsheet. Spines were classified as thin, stubby, or mushroom on the basis of the presence or absence of a clear neck region and on the basis of the head diameter. Spines lacking a discernable neck were classified as stubby. Spines with a neck region and head diameter of 0.4 µm and above were classified a mushroom, while neck containing spines with a head diameter below this value were classified as thin. Spine imaging and analysis was performed by an experimenter blinded to conditions.

#### Measurement of dendrites

DIV23 cultured rat hippocampal neurons were transfected with GFP alone or in combination with Myc-tagged mouse Rap1b. 2.5 days later cultures were fixed as described above and processed for immunohistochemistry using the same methods and antibodies as detailed in the above section. Pyramidal neuron dendrites were manually traced in Image J and dendrite length, the number of terminal dendrites, and Sholl analysis performed using Image J.

#### C-fos immunohistochemistry and quantification

Mice were allowed to explore four novel objects in a square open field box purchased from Maze Engineers (40cm long, 40cm wide, 30cm tall) for 7 minutes (mice not exposed to the novel objects were used as a control). 3 hours post novel object exposure, mice were terminally anesthetized with ketamine/xylazine and transcardially perfused for 30 seconds with PBS followed by 7 minutes 4% paraformaldehyde at a flow rate of 5-7 mls/minute. Brains were removed, post-fixed overnight at 4°C, dehydrated in 10% and 20% sucrose sequentially, and sectioned coronally on a cryostat. Hippocampal coronal sections (40 μm) were washed 3x in PBS with 0.3% Triton-X (PBS-Tx) and blocked for 1 hour at room temperature in PBS-Tx containing 3% normal donkey serum (NDS). Primary antibodies against c-Fos (Cell Signaling Cat# 2250; 1:1000) and NeuN (GeneTex Cat# GTX00837; 1:5000) were diluted in blocking solution and incubated overnight at room temperature with gentle shaking. Sections were then washed 3x in PBS and incubated for 2 hours at room temperature in a diluted mixture (1:250) containing secondary donkey-anti-rabbit-AlexaFluor-488 (Jackson Immuno; Cat# 711-545-152) and donkey-anti-chicken-AlexaFluor-594 (Jackson Immuno; Cat# 703-585-155). Unbounded antibody was removed by 3 PBS washes, and sections were mounted on slides and coverslipped with VECTASHIELD HardSet (Vector Laboratories, Burlingame, CA, USA).

The hippocampus was imaged using a Keyence BZ-X700E scanning microscope using a 10x objective lens at a 1.0 digital zoom. Z-stacks were obtained for each image using a 0.5 μm step size. Within each experiment, all microscope parameters, including digital gain and offset, were kept constant. Because the spread into the z-plane differs between individual neurons, a single image in the middle of the z-stack was chosen for analysis, where the middle is defined as the site at which the soma size is at its maximum. This approach prevents cells with greater spread into the *Z*-plane from increasing fluorescent intensity measurements. The CA3 perimeter of NeuN-positive neurons was traced and the trace was applied to the green fluorescent channel of the same *Z*-plane image containing c-Fos labeling, and the mean c-Fos fluorescent intensity within the CA3-NeuN perimeter trace was measured using ImageJ (National Institutes of Health).

The quantification of individual cell intensity and the number of cells double-labeled for c-Fos and NeuN in the dorsal CA3 region of the hippocampus was manually performed using ImageJ. A perimeter of one NeuN-positive neuron was traced and that trace was applied to each individual cell in the same *Z*-plane image containing c-Fos labeling. For each animal, the individual intensity and number of cells was averaged from 3 sections including both the left and right dorsal CA3 region of the hippocampus. All imaging and quantification was performed blind to experimental conditions.

#### Structured illumination microscopy (SIM) and associated spine nanodomain quantification

Dissociated cultures of primary hippocampal neurons were prepared from embryonic day 18 Sprague-Dawley rat embryos. Hippocampal tissue was dissected in pre-cooled HBSS that was supplemented with 1% sodium pyruvate, 0.1% D+Glucose solution (Sigma), and 10mM HEPES. Cultures were plated onto coverslips coated with poly-D-lysine (0.1mg/ml; Gibco). Trypsin solution was used for tissue dissociation in plating media (DMEM with 10% FBS, 200mM L-glutamine, and penicillin/streptomycin). 3 hours following plating, media was replaced with maintenance media (Neurobasal supplemented with B27 (Gibco), 200mM L-glutamine, and penicillin/streptomycin). Cells were treated with 1uM of Cytosine B-D-arabinofuranoside on DIV2. Beginning on DIV5 the feeding medium was supplemented with 200 μM of D,L-amino-phosphonovalerate acid (TOCRIS). 50% of the maintenance media was changed every 3-4 days. Hippocampal neurons were transfected on DIV23 using Lipofectamine 2000 with GFP and mouse Rap1b containing a Myc tag. 2.5 days post-transfection (DIV25/DIV26), neurons were fixed in 4% formaldehyde, 4% sucrose PBS for 10 minutes, followed by 10 minute fix with -20°C methanol. Coverslips were then processed for immunostaining using primary antibody against GFP (AVES, Cat# GFP-1010) and Myc (Cell Signaling, Cat# 2276).

Multichannel structured illumination microscopy (SIM) images were acquired with a Nikon Structured Illumination microscope using a 100× 1.49 NA objective and reconstructed using Nikon Elements software. z stack (z = 0.12 μm) images of secondary dendrites on pyramidal neurons were processed and analyzed using Nikon Elements software. Single spine analysis was carried out on 258 spines across 7 neurons. Imaging and reconstruction parameters were determined with the support of the Biochemistry Optical Core at the University of Wisconsin-Madison. Acquisition was set to 5MHz at 14bit with EM gain and no binning. Auto exposure was kept consistent between images and within 250-300ms, the EM gain multiplier was kept below 300, and conversion gain was held at 1x. Reconstruction parameters (Illumination Modulation Contrast, High Resolution, Noise Suppression and Out of Focus Blur Suppression) were kept consistent across all images. Maximum intensity projections were used for analysis.

Using the Nikon Elements software, dendrites and spines were outlined using binary thresholding in the channel of the cell fill, then manually corrected. Spine and neck length were manually measured, and the spine head was outlined manually. Spine classification was done using morphology, head area, and spine length. Spines lacking neck were classified as stubby. Head bearing spines with a head area of 0.4µm^2^ or larger was classified as mushroom and those with a head area below this value classified as thin. Spines with a length of 1.5µm of greater were considered filopodia. Rap1 puncta were outlined using binary thresholding and corrected manually. Visual assessment of fluorescence intensity was used to delineate connected puncta. The number of puncta within each spine and the location of the puncta was quantified manually.

Line scans were collected using the Nikon Elements Intensity profile tool. Linear sections that spanned the length of each dendritic spine were manually selected and the width was adjusted so that the selected area encompassed the entire spine. Pixel intensity graphs were created across the selected area, plotting Rap1 channel intensity against distance. The graph measurement tool was used to measure and mark the neck and head regions of the spine and the area under the curve for the Rap1 intensity graph was generated for each sub-region.

#### General behavioral procedures

All behavioral studies were performed and analyzed by an experiments who was blinded to the experimental conditions. All behaviors were recorded by an overhead digital camera with zoom function. For each behavioral test, naïve mice were used to prevent any chance of carryover effects from repeated testing.

#### Y-maze spontaneous alternation

The Y-maze procedures were adapted from our previous studies.^38,49,96^ The Y-maze was constructed by Maze Engineers and contains 3 arms (35 cm long, 5 cm wide) radiating from a central area (120⁰ separation between adjacent arms). Mice were allowed 10 minutes of free exploration of the Y-maze and the order of arm entries monitored. Arm entry criteria required that a mouse’s hind limbs were a minimum of 5 cm into an arm. The number of successive three arm alternations was logged. Proper spontaneous alternation occurs when mice enter each arm in a continual series of overlapping triplets, visiting the most recently entered arm last. The spontaneous alternation percentage was calculated as the number of overlapping arm triplet navigations divided by the total number of possible three arm entries (total arm entries subtracted by 2). As re-entry into the same arm that an animal just left is extremely rare, chance performance in this task is 50% alternation. The Y-maze was captured by an overhead digital video recorder for subsequent analysis. This task was performed and analyzed by an experimenter blinded to conditions.

#### Object-in-place testing

The object-in-place task was performed using our established procedures.^38,49^ All habituation and trials of this task were performed in a gray square open field box purchased from Maze Engineers (40cm long, 40cm wide, 30cm tall). For habituation, mice were placed in the center of the box in the absence of any novel object and allowed free navigation for 10 minutes. The next day (trial 1), four novel objects were placed about 15 cm for each corner of the open field. The objects were constructed of plastic and were about 8 cm tall. The objects all had distinct visual and textural components allow for mice to differentiate between the individual objects. During trial 1, mice were allowed free exploration of the novel object for 7 minutes. Mice were then placed into their home cage for 7 minutes, then subsequently returned to the open field for the start of trial 2. During trial 2 the same novel objects were present as during trial 1; however, the location of two the objects was swapped while the location of the other two objects was fixed between trails 1 and 2. During trial 2, mice were allowed free exploration of the objects for 7 minutes. Both trials were recorded via an overhead camera, and the video slowed to one-third real time to allow for accurate quantification. Object investigation was defined as a mouse having its nose directed within 1 cm of an object. Lying against an object was not counted as investigation. In rare instance in which a mouse attempted to climb an object, the climbing behavior was not scored as object investigation. Time spent in direct object investigation was quantified using a digital stopwatch with a certified 0.01 second resolution. The recognition ratio was calculated as the amount of time mice spent investigating the swapped object pair divided by the time spent investigating the non-swapped pair, with chance performance level=1.0. This task was performed and quantified by an experimenter blinded to conditions.

#### Quantification and statistical analysis

Graphpad Prism version 9 was used for all statistical analyses. Comparison between two groups was performed using a Student’s two-tailed t-test when groups followed a normal distribution and had normal variance. When variance between groups differed substantially, a Welch’s t-test was used, and when data did not follow a normal distribution, Mann Whitney was used. Comparisons between four conditions across two sets of variables was performed using a two-way ANOVA with a Bonferroni post hoc for multiple comparison correction. Object-in-place analysis across the five successive time bins was performed using a repeated measures two-way ANOVA with a Holm-Sidak post hoc for multiple comparison correction. A standard one-way ANOVA was used when comparing three or more groups along a single parameter, with Bonferroni post hoc for multiple comparison, except if the data were not normally distributed in which case a Welch’s one-way ANOVA and Dunnett’s post hoc was used. Grubb’s test was used for potential outlier detection in the biochemical data, while the ROUT method was used for potential outlier detection of the SIM data. Spine head diameter survival curves were analyzed using a log-rank (Mantel-Cox test). Assessment for normal data distribution were performed using the Shapiro-Wilk test and assessment for normal variance performed using F-tests. On graphs, *p<0.05, **p<0.01, ***p<0.001. The results of statistical analyses are reported in the main text, while the group sample sizes are reported in the figure legends.

