## Supplemental Figures and Figure Legends for "Aberrant regulation of the Rap1 small GTPase in response to escalating, intermittent stress produces hippocampal synaptic and cognitive dysfunction"

**Figure S1**

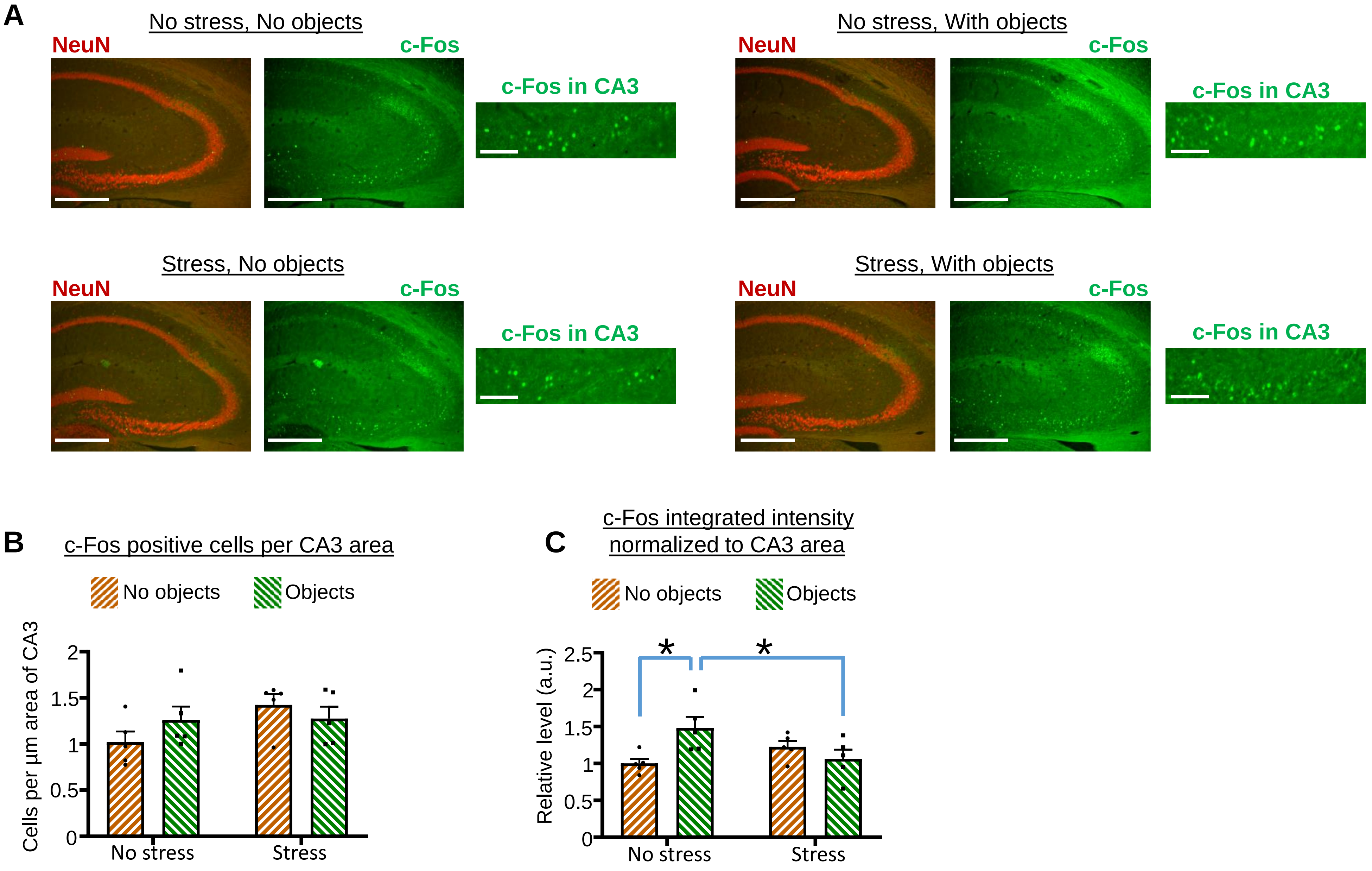
A
No stress, No objects
NeuN
c-Fos
c-Fos in CA3
No stress, With objects
NeuN
c-Fos
c-Fos in CA3
Stress, No objects
NeuN
c-Fos
c-Fos in CA3
Stress, With objects
NeuN
c-Fos
c-Fos in CA3
B
c-Fos positive cells per CA3 area
No objects
Objects
Cells per µm area of CA3
0
0.5
1
1.5
2
No stress
Stress
C
c-Fos integrated intensity normalized to CA3 area
No objects
Objects
Relative level (a.u.)
0
0.5
1
1.5
2
2.5
*
*
No stress
Stress

#### Figure S2

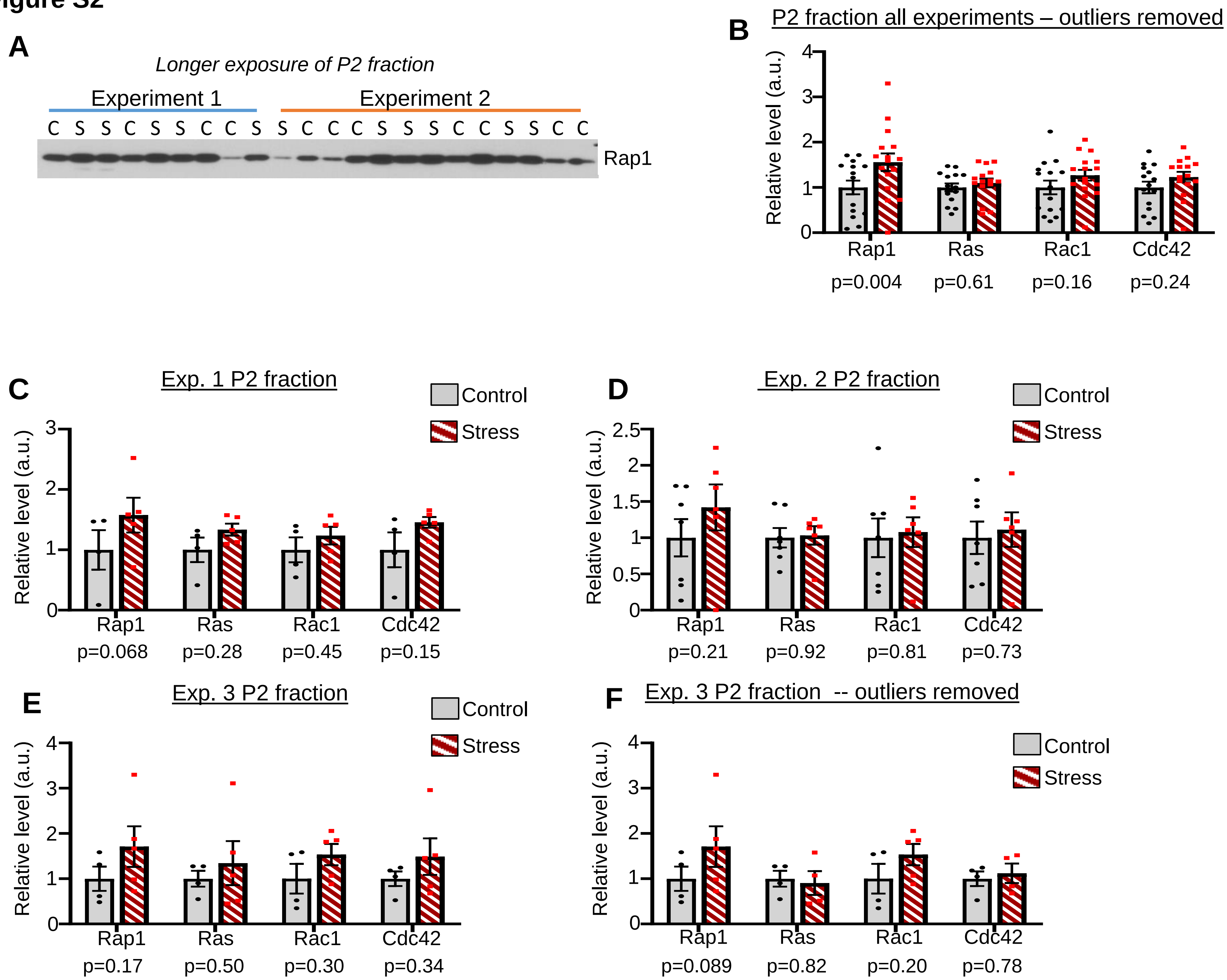
A
Longer exposure of P2 fraction
Experiment 1
Experiment 2
C S S C S S C C S S C C C S S S C C S S C C
Rap1
B
P2 fraction all experiments – outliers removed
Relative level (a.u.)
0
1
2
3
4
Rap1
Ras
Rac1
Cdc42
p=0.004
p=0.61
p=0.16
p=0.24
C
Exp. 1 P2 fraction
Control
Stress
Relative level (a.u.)
0
1
2
3
Rap1
Ras
Rac1
Cdc42
p=0.068
p=0.28
p=0.45
p=0.15
D
Exp. 2 P2 fraction
Control
Stress
Relative level (a.u.)
0
0.5
1
1.5
2
2.5
Rap1
Ras
Rac1
Cdc42
p=0.21
p=0.92
p=0.81
p=0.73
E
Exp. 3 P2 fraction
Control
Stress
Relative level (a.u.)
0
1
2
3
4
Rap1
Ras
Rac1
Cdc42
p=0.17
p=0.50
p=0.30
p=0.34
F
Exp. 3 P2 fraction -- outliers removed
Control
Stress
Relative level (a.u.)
0
1
2
3
4
Rap1
Ras
Rac1
Cdc42
p=0.089
p=0.82
p=0.20
p=0.78

**Figure S3**

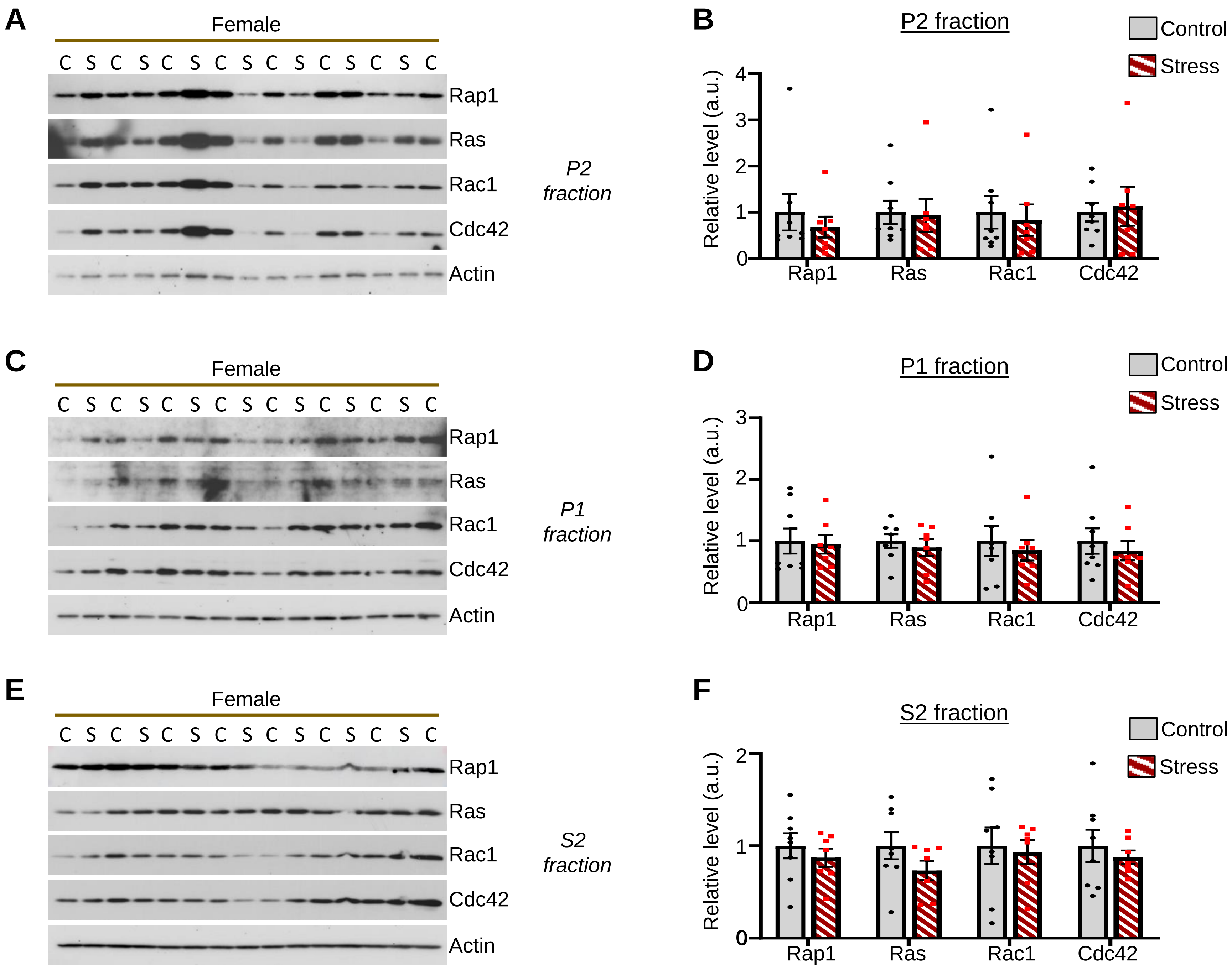
A
Female
C S C S C S C S C S C S C S C
Rap1
Ras
Rac1
Cdc42
Actin
P2 fraction
B
P2 fraction
Control
Stress
Relative level (a.u.)
0
1
2
3
4
Rap1
Ras
Rac1
Cdc42
C
Female
C S C S C S C S C S C S C S C
Rap1
Ras
Rac1
Cdc42
Actin
P1 fraction
D
P1 fraction
Control
Stress
Relative level (a.u.)
0
1
2
3
Rap1
Ras
Rac1
Cdc42
E
Female
C S C S C S C S C S C S C S C
Rap1
Ras
Rac1
Cdc42
Actin
S2 fraction
F
S2 fraction
Control
Stress
Relative level (a.u.)
0
1
2
Rap1
Ras
Rac1
Cdc42

**Figure S4**

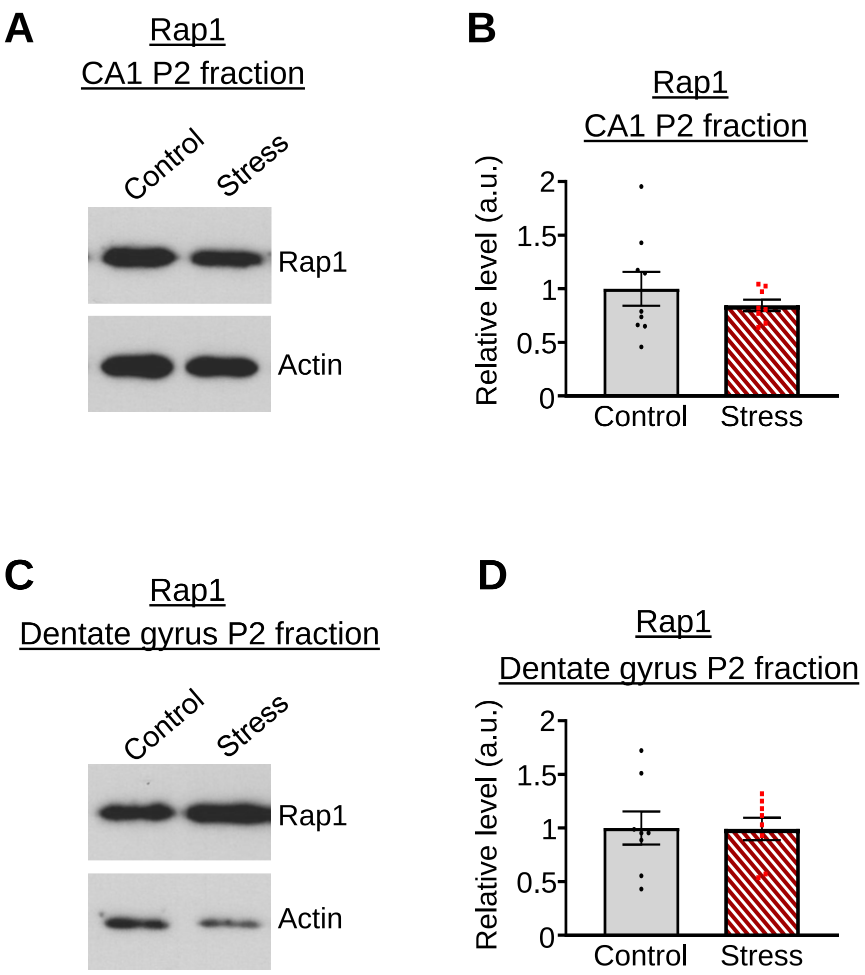
A
Rap1
CA1 P2 fraction
Control
Stress
Rap1
Actin
B
Rap1
CA1 P2 fraction
Relative level (a.u.)
2
1.5
1
0.5
0
Control
Stress
C
Rap1
Dentate gyrus P2 fraction
Control
Stress
Rap1
Actin
D
Rap1
Dentate gyrus P2 fraction
Relative level (a.u.)
2
1.5
1
0.5
0
Control
Stress

#### Figure S5

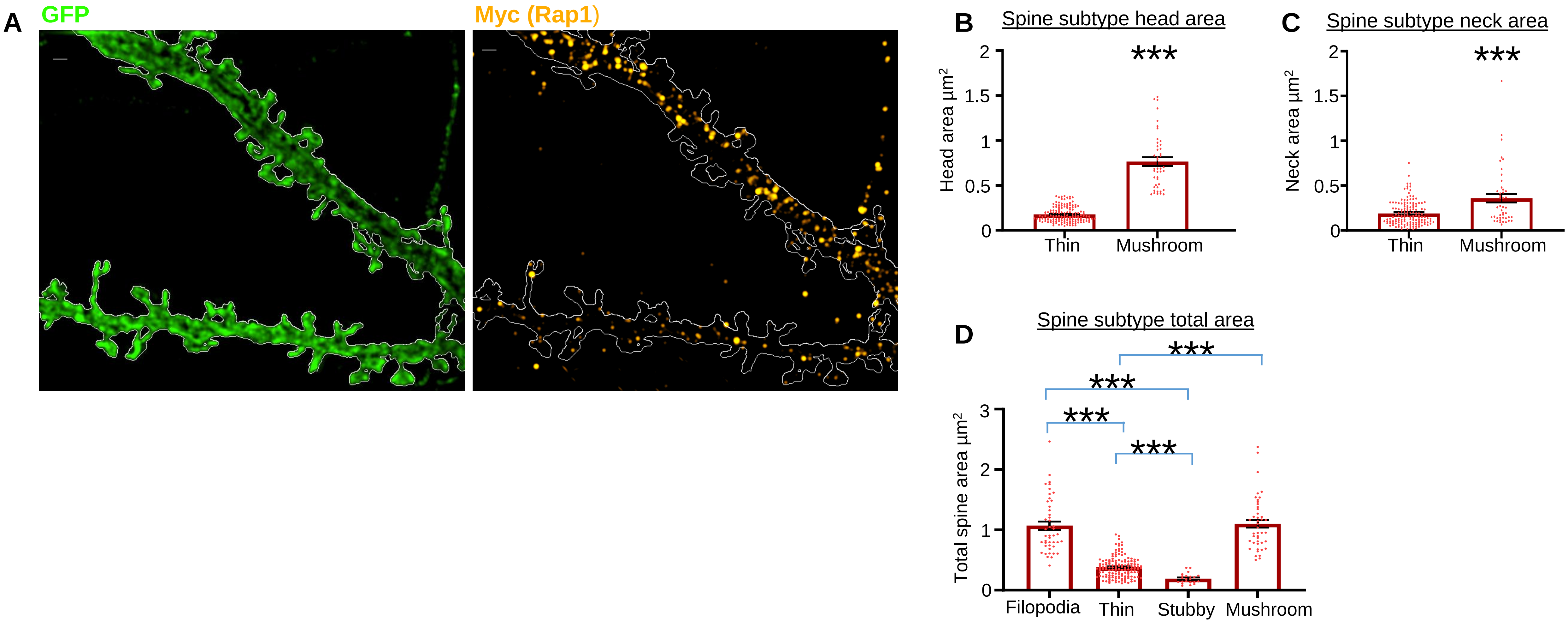
A
GFP
Myc (Rap1)
B
Spine subtype head area
Head area µm²
2
1.5
1
0.5
0
***
Thin
Mushroom
C
Spine subtype neck area
Neck area µm²
2
1.5
1
0.5
0
***
Thin
Mushroom
D
Spine subtype total area
Total spine area µm²
3
2
1
0
***
***
***
***
Filopodia
Thin
Stubby
Mushroom

#### Figure S6

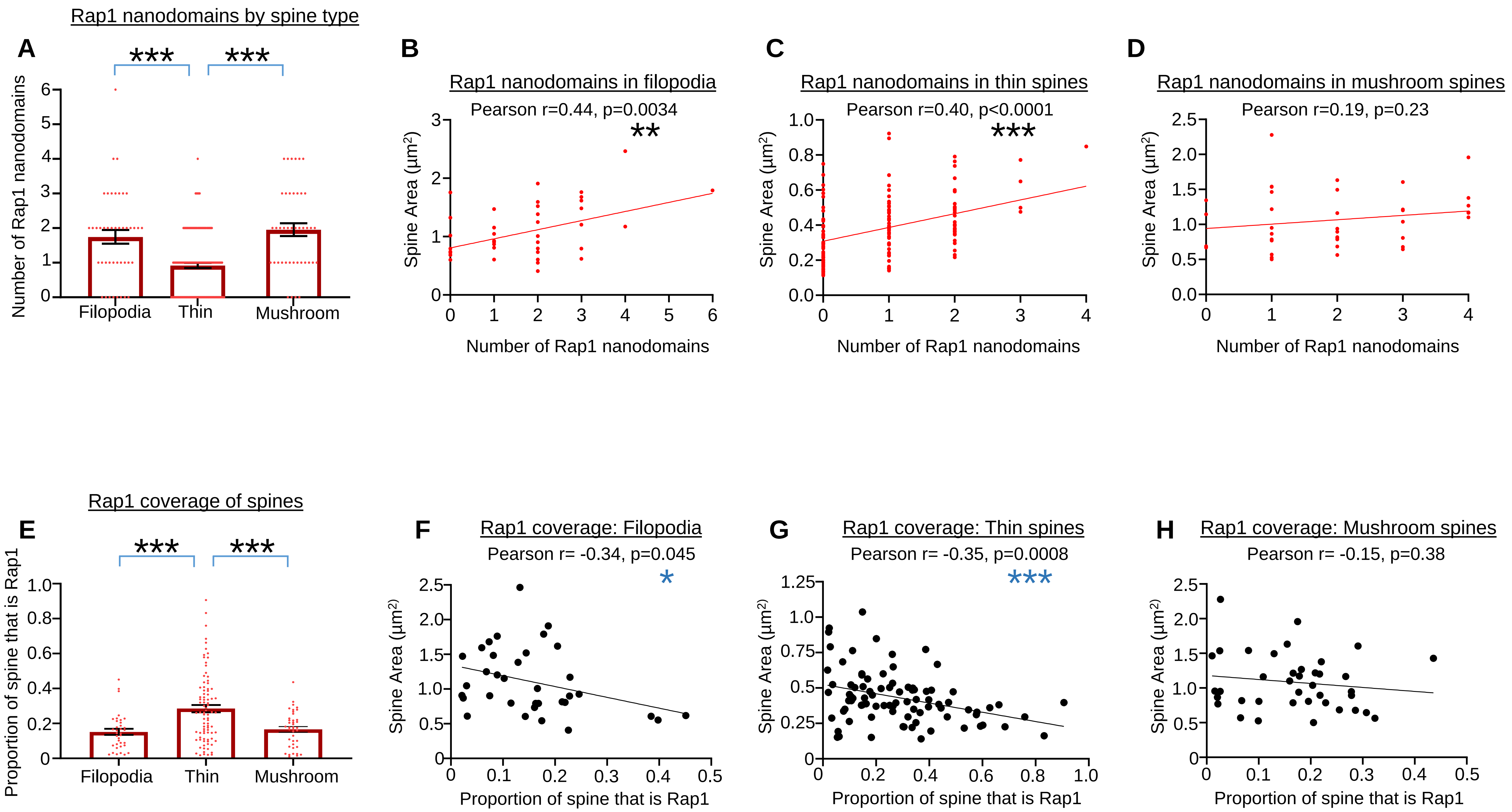

Rap1 nanodomains by spine type
A
***
***
Number of Rap1 nanodomains
0
1
2
3
4
5
6
Filopodia
Thin
Mushroom
B
Rap1 nanodomains in filopodia
Pearson r=0.44, p=0.0034
**
Spine Area (µm2)
Number of Rap1 nanodomains
C
Rap1 nanodomains in thin spines
Pearson r=0.40, p<0.0001
***
D
Rap1 nanodomains in mushroom spines
Pearson r=0.19, p=0.23
Rap1 coverage of spines
E
***
***
Proportion of spine that is Rap1
F
Rap1 coverage: Filopodia
Pearson r= -0.34, p=0.045
*
G
Rap1 coverage: Thin spines
Pearson r= -0.35, p=0.0008
***
H
Rap1 coverage: Mushroom spines
Pearson r= -0.15, p=0.38

### Figure S7

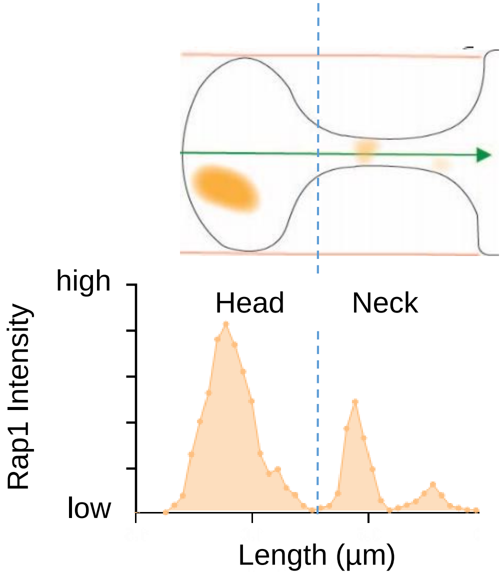

Figure S8

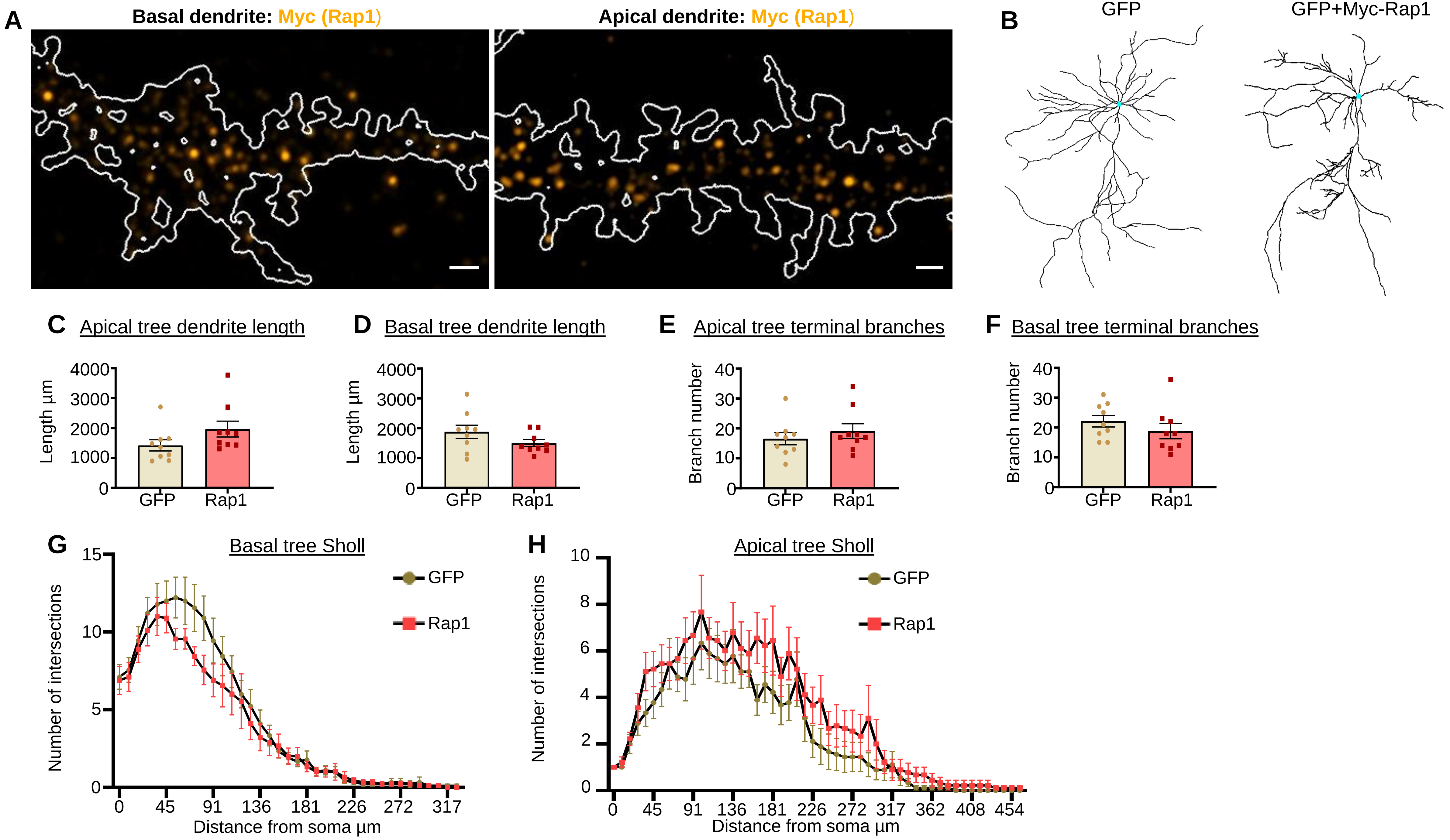

A
Basal dendrite: Myc (Rap1)
Apical dendrite: Myc (Rap1)
B
GFP
GFP+Myc-Rap1
C Apical tree dendrite length
Length µm
4000
3000
2000
1000
0
GFP
Rap1
D Basal tree dendrite length
Length µm
E Apical tree terminal branches
Branch number
40
30
20
10
0
F Basal tree terminal branches
G Basal tree Sholl
GFP
Rap1
Number of intersections
15
10
5
0
0 45 91 136 181 226 272 317
Distance from soma µm
H Apical tree Sholl
10
8
6
4
2
0
0 45 91 136 181 226 272 317 362 408 454

**Figure S9**

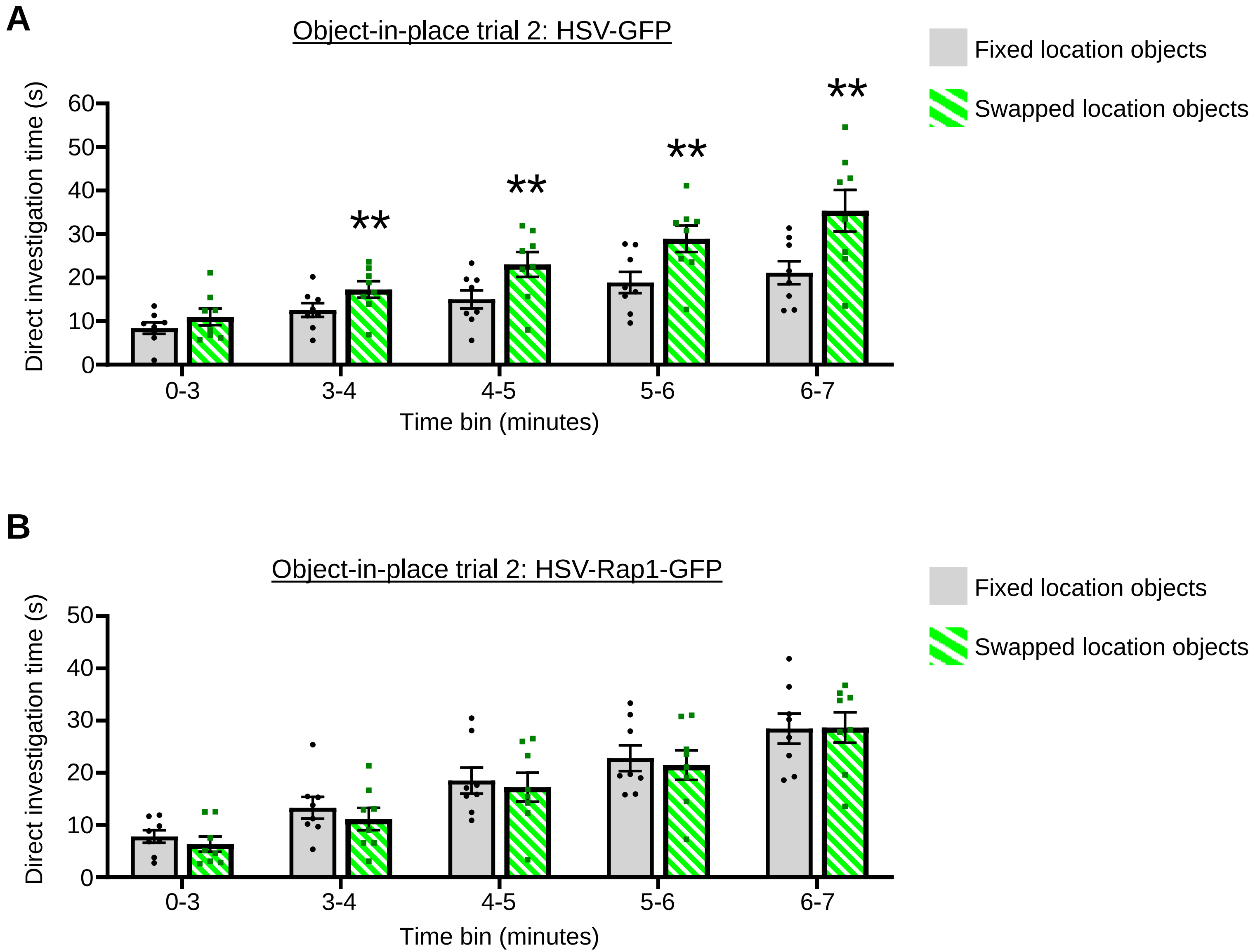
A
Object-in-place trial 2: HSV-GFP
Fixed location objects
Swapped location objects
Direct investigation time (s)
60
50
40
30
20
10
0
**
**
**
**
0-3
3-4
4-5
5-6
6-7
Time bin (minutes)
B
Object-in-place trial 2: HSV-Rap1-GFP
Fixed location objects
Swapped location objects
Direct investigation time (s)
50
40
30
20
10
0
0-3
3-4
4-5
5-6
6-7
Time bin (minutes)

**Supplementary Figure legends**

Figure S1. Additional images and analysis of escalating, intermittent stress on c-Fos immuno-reactivity in the CA3 region.

A. Additional low magnification images of NeuN and c-Fos immuno-label in the hippocampus (with CA3 region visible) of non-stressed and stressed mice that were either not exposed to novel objects or were allowed to explore novel objects (Scale bar=300μm). High magnification images exclusively show the CA3 region and the associated c-Fos immuno-label (Scale bar=10μm).

B. Graph depicts the number of c-Fos positive cells within the CA3 region in each stress and object exploration condition normalized to the area of the CA3 measured. No significant main effects, interactions, or differences between groups were detected. Cell count is the combined total from three coronal planes of the CA3. n=5 no stress, no object mice; 5 no stress, object mice; 5 stress, no object mice; 5 stress, object mice.

C. Graph depicts c-Fos integrated intensity in each stress and object exploration condition normalized to the area of the CA3 measured. A significant interaction between stress status and object investigation status was identified [$F(1,16)=8.767$, $p=0.0092$].*Bonferroni post hoc between indicated comparisons, $p<0.05$. The integrated intensity of all cells within a single mouse were averaged, and this value used for statistical analyses. n=5 no stress, no object mice; 5 no stress, object mice; 5 stress, no object mice; 5 stress, object mice.

All summary data are the mean + SEM.

Figure S2. Effects of escalating, intermittent stress on small GTPases levels in CA3 field P2 fraction within individual experiments.

A. Long Western blot exposure of Rap1 in CA3 region P2 fraction from experiment 1 and experiment 2. Lighter exposure is shown in the main figure.

B. Graph depicts quantification of the indicated small GTPases in the CA3 region P2 homogenates from non-stressed and stressed mice with outliers removed (Grubb's test, 1 outlier in stress Ras, and 1 outlier in stress Cdc42). n=15 non-stressed and 16 stressed mice. Direct comparison p values (Fisher's LSD) are indicated below the graph for each small GTPase.

C-F. Graphs depict small GTPases levels in CA3 fractions from control and stress mice within individual experiments (the combined analysis across all experiments is shown in the main figure). Fisher's LSD direct comparison p-values for each small GTPase are indicated. Note that for experiment 3 a single stress sample was a Grubbs' test outlier for Ras and Cdc42. The p values with and without this outlier are shown for experiment 3 in S2E and S2F, respectively.

All summary data are the mean + SEM.

Figure S3. Effects of escalating, intermittent stress on small GTPase expression profiles in female mice.

A. P2 fractions derived from CA3 homogenates of non-stressed and stressed female mice. Blots were probed for the indicated small GTPases. C=non-stressed controls, S=stressed.

B. Stress did not affect levels of Rap1, Ras, Rac1, or Cdc42 in female CA3 P2 fractions. Bonferroni $p>0.9999$ for all four small GTPases. n=8 control and 7 stressed mice.

C. P1 fractions derived from CA3 homogenates of non-stressed and stressed female mice. Blots were probed for the indicated small GTPases. C=non-stressed controls, S=stressed.

D. Stress did not affect levels of Rap1, Ras, Rac1, or Cdc42 in female CA3 P1 fractions. Bonferroni $p>0.9999$ for all four small GTPases. n=8 control and 7 stressed mice.

E. S2 fractions derived from CA3 homogenates of non-stressed and stressed female mice. Blots were probed for the indicated small GTPases. C=non-stressed controls, S=stressed.

F. Stress did not affect levels of Rap1 (Bonferroni $p>0.9999$), Ras (Bonferroni $p=0.7624$), Rac1 (Bonferroni $p>0.9999$), or Cdc42 (Bonferroni $p>0.9999$) in female CA3 S2 fractions. n=8 control and 7 stressed mice.

All summary data are the mean + SEM.

Figure S4. Effects of scalating intermittent stress on Rap1 levels in the CA1 hippocampal field and dentate gyrus.

A. P2 fractions derived from CA1 homogenates of non-stressed mice and mice subjected to escalating, intermittent stress (tissue collected 24 hour after final stress exposure). Blots were probed for Rap1.

B. Stress does not affect Rap1 levels in CA1 P2 fractions [$t(15)=0.8831$, $p=0.3911$]. n=9 non-stressed and 8 stressed mice.

C. P2 fractions derived from dentate gyrus homogenates of non-stressed mice and mice subjected to escalating, intermittent stress (tissue collected 24 hour after final stress exposure). Blots were probed for Rap1.

D. Stress does not affect Rap1 levels in dentate gyrus P2 fractions [t(14)=0.04330, p=0.9661]. n=8 non-stressed and 8 stressed mice.

All summary data are the mean + SEM.

Figure S5. Dendritic spine measurement parameters among protrusions imaged via SIM.

A. Low magnification image showing overexpressed Myc-tagged Rap1 nanodomains (labeled via anti-Myc tag antibody) in mature cultured hippocampal neurons.

B. Mushroom spines have a greater head area than thin spines. Unpaired t-test, [t(184)=20.19, p<0.0001]. n=143 thin spines, 43 mushroom spines.

C. Mushroom spines have a greater neck area than thin spines. Unpaired t-test, [t(186)=5.048, p<0.0001]. n=143 thin spines, 45 mushroom spines.

D. Graph depicts total spine area for filopodia, thin spines, stubby spines, and mushroom spines. Differences between groups are indicated on the graph. One-way ANOVA with Dunnett's multiple comparison correction. ***p<0.0001. n=45 filopodia, 142 thin spines, 22 stubby spines, 44 mushroom spines

All summary data are the mean + SEM.

Figure S6. Myc-Rap1 nanodomains and Myc-Rap1 coverage of different protrusion sub-types.

A. Graph depicts the mean number of Myc-Rap1 nanodomains for filopodia, thin spines, and mushroom spines. Thin spines show a lower number of nanodomains than either filopodia or mushroom spines. One way ANOVA F(2,224)=0.4.025, p=0.0192; Bonferroni post hoc comparisons filopodia vs. thin, p<0.0001; filopodia vs. mushroom, p>0.9999; thin vs. mushroom, p<0.0001. n=43 filopodia, 142 thin, 42 mushroom; from 7 neurons.

B. Graph depicts the impact of spine area on the number of Myc-Rap1 nanodomains in filopodia. A significant positive correlation was detected. Pearson r=0.44, p=0.0034. n=43 filopodia; from 7 neurons

C. Graph depicts the impact of spine area on the number of Myc-Rap1 nanodomains in thin spines. A significant positive correlation was detected. Pearson r=0.40, p<0.0001. n=142 thin spines; from 7 neurons

D. Graph depicts the impact of spine area on the number of Myc-Rap1 nanodomains in mushroom spines. No significant correlation was detected. Pearson r=0.19, p=0.23. n=42 mushroom spines; from 7 neurons

E. Graph depicts the proportion of the indicated protrusion areas occupied by Myc-Rap1. Thin spines show a greater proportion of their area occupied by Myc-Rap1 compared to filopodia or mushroom spines. One way ANOVA F(2,160)=11.46, p<0.0001; Bonferroni post hoc comparisons filopodia vs. thin, p=0.0002; filopodia vs. mushroom, p>0.9999; thin vs. mushroom, p=0.0005. n=36 filopodia, 87 thin, 40 mushroom; from 7 neurons.

F. Graph depicts the impact of spine area on the proportion of the filopodia that is occupied by Myc-Rap1. A significant negative correlation was detected. Pearson r= -0.3409, p=0.045. n=36 filopodia; from 7 neurons

G. Graph depicts the impact of spine area on the proportion of a thin spine that is occupied by Myc-Rap1. A significant negative correlation was detected. Pearson r= -0.3541, p=0.0008. n=87 thin spines; from 7 neurons.

H. Graph depicts the impact of spine area on the proportion of a mushroom spine that is occupied by Myc-Rap1. No significant correlation was detected. Pearson r= -0.15, p=0.38. n=40 mushroom spines; from 7 neurons.

All summary data are the mean + SEM.

Figure S7. Depiction of line scan of Rap1 intensity through the head and neck of a spine.

Line scan begins at the apex of the head region and continues to the point where the spine neck joins the underlying dendrite shaft. The width of the scan region was manually adjusted for each spine to assure it captures the entirety of the spine head. Delineation is made between the Rap1 intensity in the head and that of the neck for every analyzed spine, and intensity values are background subtracted. On the image, the green arrow depicts the direction of the line scan, and the orange lines depict the width of the line scan.

Figure S8. The effects of Rap1 overexpression on cultured hippocampal dendrite morphology.

A. DIV23 cultured hippocampal neurons were transfected with GFP alone or GFP in combination with Myc-tagged Rap1. Immunohistochemistry was used to label the Myc tag and SIM reveals the presence of Rap1 nanodomains throughout the dendrite shaft of apical and basal dendrites. Scale bar=1 μm

B. Representative dendrite tracings of cultured hippocampal neurons expressing GFP alone or GFP in combination with Myc-tagged Rap1.

C. Rap1 overexpression did not affect total dendrite length of the apical tree [t(16)=1.685, p=0.1114]; n=9 GFP neurons, 9 GFP+Rap1 neurons.

D. Rap1 overexpression did not affect total dendrite length of the basal tree [t(16)=1.530, p=0.1455]; n=9 GFP neurons, 9 GFP+Rap1 neurons.

E. Rap1 overexpression did not affect total number of terminal dendrites of the apical tree [t(16)=0.8031, p=0.4337]; n=9 GFP neurons, 9 GFP+Rap1 neurons.

F. Rap1 overexpression did not affect total number of terminal dendrites of the basal tree [t(16)=1.039, p=0.3141; n=9 GFP neurons, 9 GFP+Rap1 neurons.

G. Sholl analysis indicates that Rap1 overexpression did not affect the branching complexity of the basal dendrite tree either globally [Rap1 main effect F(1,16)=1.594, p=0.2248], or at any intersections from the soma (Fisher's LSD, p>0.05 at all intersections). n=9 GFP neurons, 9 GFP+Rap1 neurons.

H. Sholl analysis indicates that Rap1 overexpression did not affect the branching complexity of the apical dendrite tree either globally [Rap1 main effect F(1,16)=3.060, p=0.0994], or at any intersections from the soma (Fisher's LSD, p>0.05 at all intersections). n=9 GFP, 9 GFP+Rap1 neurons.

All summary data are the mean + SEM.

Figure S9. Raw investigation times of object-in-place performance.

A. Graph depicts the amount of time mice overexpressing HSV-GFP into the CA3 region investigate object pairs during trial 2 of the object-in-place task. Time investigating the fixed location objects and the objects that swapped location between trials is shown. HSV-GFP mice spent more time investigating the swapped objects pair vs. the fixed object location pair at all time bins except the 0-3 bin. Sidak multiple comparison corrected p-vales 0-3 minute bin (p=0.2491), 3-4 minute bin (p=0.0031), 4-5 minute bin (p=0.0031), 5-6 minute bin (p=0.0016), 6-7 minute bin (p=0.0031).

B. Graph depicts the amount of time mice overexpressing HSV-Rap1-GFP into the CA3 region investigate object pairs during trial 2 of the object-in-place task. Time investigating the fixed location objects and the objects that swapped location between trials is shown. No difference in the amount of time spent investigating the swapped objects vs. the fixed location objects was detected. Sidak multiple comparison corrected p-vales 0-3 minute bin (p=0.4078), 3-4 minute bin (p=0.8505), 4-5 minute bin (p=0.9425), 5-6 minute bin (p=0.9425), 6-7 minute bin (p=0.9425).

All summary data are the mean + SEM.
